# *Himito*: a Graph-based Toolkit for Mitochondrial Genome Analysis using Long Reads

**DOI:** 10.1101/2025.11.03.686348

**Authors:** Hang Su, Yongqing Huang, Timothy Durham, Nahyun Kong, Emma Casey, David Benjamin, Sheng Chih Jin, Kiran V Garimella

**Affiliations:** Data Sciences Platform, Broad Institute of MIT and Harvard, Cambridge, MA, USA; Metabolism Program, Broad Institute of MIT and Harvard, Cambridge, MA, USA; Department of Genetics, Washington University School of Medicine, St. Louis, MO 63110, 11 USA

## Abstract

Understanding the genetic and epigenetic regulation of mitochondrial DNA (mtDNA) is essential for elucidating mechanisms of aging and disease. Long-read sequencing can span the entire mitochondrial genome and directly capture base-modification signals, yet analytical tools for such data remain limited. We developed *Himito*, a graph-based toolkit for analyzing mitochondrial genome using long reads. *Himito* filters reads originating from nuclear mitochondrial insertions (NUMTs), constructs a sequence graph to represent mtDNA diversity, assembles primary haplotypes, calls variants, and analyzes 5-methylcytosine (5mC) modifications within a unified framework. Benchmarking on high-quality reference datasets shows *Himito* achieves superior performance in assembly and variant calling compared with existing tools. Applied to the *All of Us* (AoU) v8 dataset, *Himito* identified pathogenic mtDNA variants, revealed population-scale haplogroup diversity, and uncovered age-related genetic and epigenetic patterns. These results demonstrate that long-read sequencing, combined with graph-based analysis, enables integrated characterization of mitochondrial genomic and epigenomic variation. *Himito* is available at https://github.com/broadinstitute/Himito.

## Introduction

Mitochondria are essential organelles responsible for cellular energy production, metabolic regulation, and aging^1,2^. Human mitochondrial DNA (mtDNA) is a 16 kb circular, double-stranded DNA molecule distinguished by its high copy number, elevated mutation rate relative to nuclear genomes, maternal inheritance, and tissue-specificity^3,4^. Mitochondrial variants contribute to diverse disorders, from metabolic and neurodegenerative diseases to cancer^5^. These variants occur as homoplasmies, shared by all the mtDNA copies within a cell, or heteroplasmies, representing mixtures of mtDNA types. Characterizing the genetic variations in mtDNA is essential for decoding the mitochondrial dysfunction and related mitochondrial diseases. These variants include single-nucleotide variants (SNVs), small insertions or deletions (indels), and large structural variants. To date, over a hundred high-confident pathogenic variants have been reported^6^, and the majority of these variants cause disease in the heteroplasmic state, when the level of mtDNA with these variants exceeds a disease-specific threshold. Thus, sensitive and accurate detection of heteroplasmy is crucial for understanding, accurate diagnosis and early intervention of mitochondrial disease^7^.

Accurate mitochondrial genome analysis is complicated by several biological and technical factors: the circular nature, the presence of heteroplasmy, and the confounding factor of nuclear-mitochondrial segments (NUMTs) – fragments of mtDNA inserted into the nuclear genome^8^. The nuclear genome harbors hundreds to thousands of ancient NUMTs^9^, introducing complexity to variant calling and heteroplasmy analysis. NUMTs insertion is also an ongoing process^10^, occurring approximately once in every 10,000 births^8^. Short reads struggle to distinguish NUMTs-derived from true mitochondrial reads, generating false-positive variant calls and spurious heteroplasmy. Moreover, mtDNA harbors large structural variants^11^ and variable homopolymer tracts that are poorly resolved by short reads^12^. Another persistent challenge is the controversial existence of 5-methylcytosine (5mC) modifications in mtDNA. Although several studies suggest report negligible or artifactual mtDNA methylation^13–15^, others report the low-level 5mC^16–18^ and mitochondrial localization of DNA methyltransferases (DNMT1, DNMT3A, DNMT3B)^19–21^,suggesting potential regulatory roles^22–24^. These conflicting observations may stem from methodological artifacts in bisulfite sequencing ^25^, including incomplete conversion of super-coiled mtDNA and mis-mapping of methylated NUMT reads^1426^. Long read sequencing circumvents these limitations by generating native reads that preserve methylation information and span the entire mitochondrial genome without PCR amplification.

Despite extensive development of long-read bioinformatics tools (https://long-read-tools.org/), most are primarily designed for diploid nuclear genomes rather than organelle-specific analysis. Common variant calling tools, such as DeepVariant^27^, are not specifically designed for mtDNA variant calling and have limited power detecting heteroplasmic variants. Tools such as *GATK-Mutect2* mitochondrial mode^28^, *mtDNA-Server*^29^, *MToolBox*^30^ are designed for short read sequence analysis, but are not optimized for long reads. *MitoHiFi*^31^ has been developed for mitochondrial genome assembly using HiFi long reads but does not perform variant calling. *Mitorsaw* (unpublished) primarily focuses on mitochondrial variant calling using PacBio HiFi long reads, but has not been tested thoroughly in literature. Consequently, no existing methods integrate NUMT filtering, haplotype-resolved assembly, variant calling, and methylation profiling of mtDNA from long reads within a unified framework.

To address this gap, we present *Himito*, a graph-based toolkit designed for mitochondrial genome analysis using long-read sequencing. *Himito* classifies and removes NUMT-derived reads based on alignment signatures and methylation patterns, constructs sequence graphs capturing the circular topology of mtDNA, assembles haplotypes, detects both homoplasmic and heteroplasmic variants, and annotates methylation events on the resulting graph. Benchmarking across multiple reference datasets demonstrates that *Himito* improves mitochondrial assembly accuracy and heteroplasmic variant detection compared with existing approaches. Application to the All of Us (AoU) v8 long-read cohort further reveals population-scale mtDNA diversity, pathogenic variants, and age-related genetic and epigenetic changes, underscoring the power of integrating graph-based and methylation-aware analysis for mitochondrial genomics.

## Results

### *Himito* algorithm

*Himito* incorporates 5 key modules for long-read-based mitochondrial genome analysis (Figure 1): `Filter, Build, Asm, Call, and Methyl`. *Himito* `Filter` module excludes the NUMTs contamination by identifying NUMTs-derived reads based on the whole-genome supplementary alignment signature (SA tags) and read-level methylation patterns (Figure 1a). *Himito* `Build` module constructs a sequence graph by mapping unique kmers in the standard human reference mitochondrial genome (rCRS), deemed as *anchors*, to the mtDNA-derived long reads (Figure 1b). The *Himito* graph captures the circular nature of mtDNAs. The local topology of the graph reveals the sequence diversity within a genomic region. The graphical genomes are stored in the standard *GFA* file format with annotations in the JSON format. The *Himito* `Asm` module extracts the primary haplotype by traversing the graph following the most-supported paths (Figure 1c). The primary haplotypes of each sample are stored in the standard FASTA files. *Himito* `Call` module aims to call both homoplasmic and heteroplasmic variants from the graph. Pair-wise alignment is performed between reference paths and alternative paths separated by anchor pairs (Figure 1d). Genetic variants are recorded on the graph in the form of Concise Idiosyncratic Gapped Alignment Report (CIGAR) strings and are mapped back to the reference coordinates. To distinguish true variants from sequence artifacts, a permutation test was employed, leveraging the variants’ phasing information to exclude false positive calls (Methods). The variants are reported in a standard biallelic VCF file with annotations. *Himito* `Methyl` module annotates the methylation signals from the graph and aggregates the methylation signals along the major haplotype. The methylation modification scores on the major haplotype are recorded in a standard BED file (Figure 1e).

**Figure 1.**
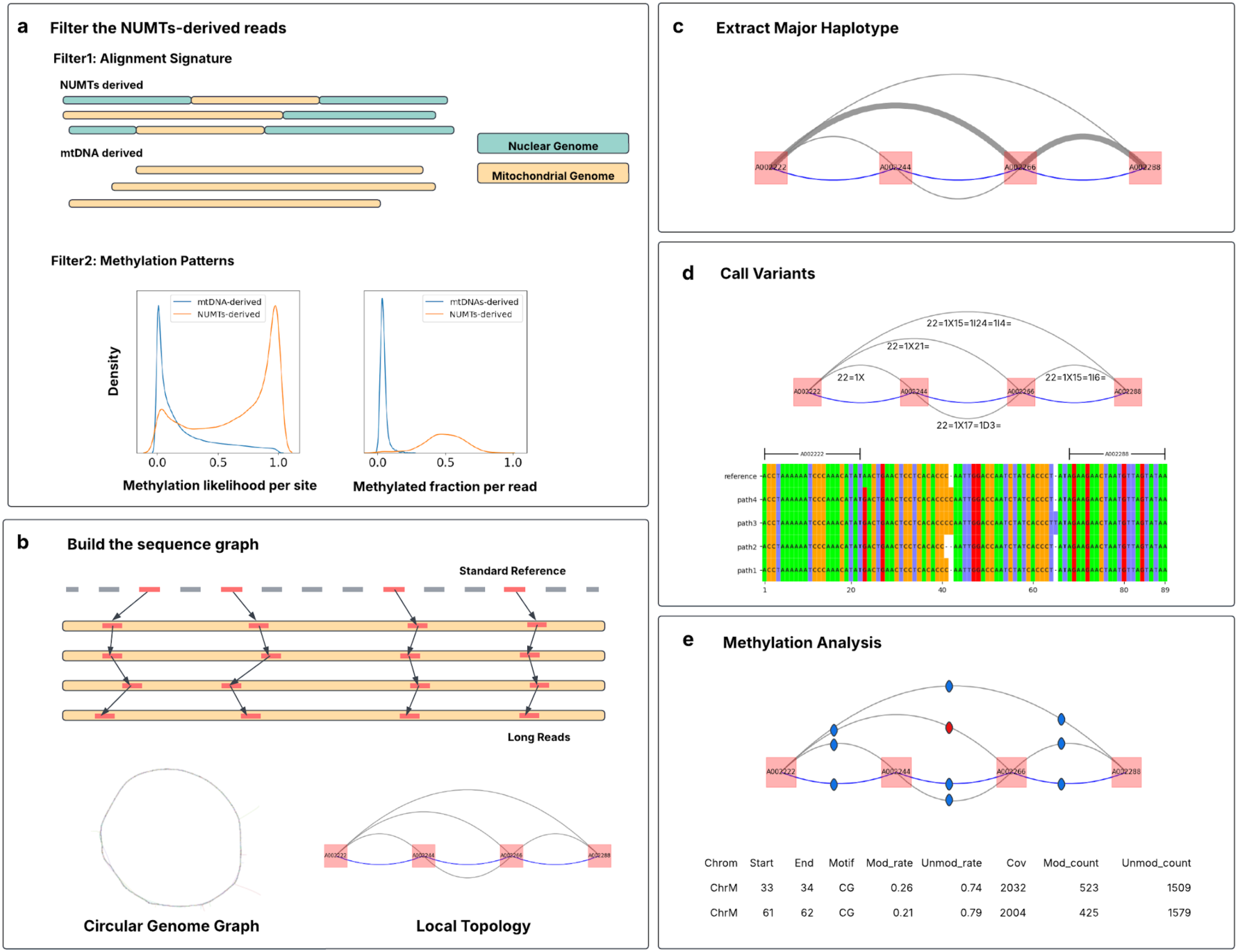
An overview of *Himito*. *Himito* comprises 5 modules: `Filter, Build, Asm, Call` and `Methyl`. a. *Himito* classifies NUMTs-derived reads and mtDNA-derived reads based on alignment signature and methylation patterns. b. The long reads identified as originating from mtDNAs are utilized to construct a sequence graph. Unique kmers (red) in the reference genome are selected as anchors and compressed the reads into a graphical genome. The graph automatically forms a circular genome, which reveals the circular nature of reads derived from mtDNAs. Within a specific genomic region, the local graph topology illustrates sequence diversity. Standard reference genomes are represented as blue paths between anchor pairs, while alternative paths, shown in grey, incorporate all sequence variations. c. Traversing the graph by following the most supported path (grey path, wide), the major haplotype of each sample is constructed. d. Sequence variants are annotated to the graph in forms of cigar strings and are further reported in standard VCF files. e. 5-methylcytosine (5mC) modification sites with their methylation likelihood are annotated to the graph and further aggregated in a standard BED file.

The innovation of *Himito* lies in its ability to accurately distinguish true mtDNA-derived reads from NUMT contaminants and to represent mitochondrial diversity through a sequence graph. *Himito* leverages both long-read sequence alignment and methylation features to filter out NUMTs-derived reads, integrating both genetic and epigenetic variation into a unified graph representation with haplotype information. The graph topology captures the circular nature of mtDNAs and delineates conserved versus highly variable regions. *Himito* also ensures compatibility with other bioinformatic pipelines by using common file formats. *Himito* is written in Rust, and serves as a fast and comprehensive toolkit for mitochondrial genetic and epigenetic analysis using long reads.

### Benchmark

We benchmarked *Himito’s* performance using PacBio long-read whole-genome sequencing (lrWGS) data from the Human Pangenome Reference Consortium (HPRC)^32^ and the HapMap mixture benchmarking dataset released in Somatic Mosaicism across Human Tissues consortium (SMaHT)^33^. The HPRC year 1 dataset includes 47 samples, each paired with high-quality haplotype-resolved whole genome assemblies.

We first compared *Himito’s* major haplotype assemblies to those generated by *MitoHifi*^31^ using the 47 HPRC samples, with the high-quality HPRC assemblies serving as the truth set. *MitoHiFi* frequently failed when the sequence depth was low (<20x). For successful runs, we computed the Jaccard similarity between the assembled mitochondrial genomes and truth assemblies by breaking the sequences into 100-, 300-, and 500-mer windows (Supplementary Methods). *Himito* produced more accurate assemblies than *MitoHiFi* across all depths, particularly below 50x (Figure 2a).

**Figure 2.**
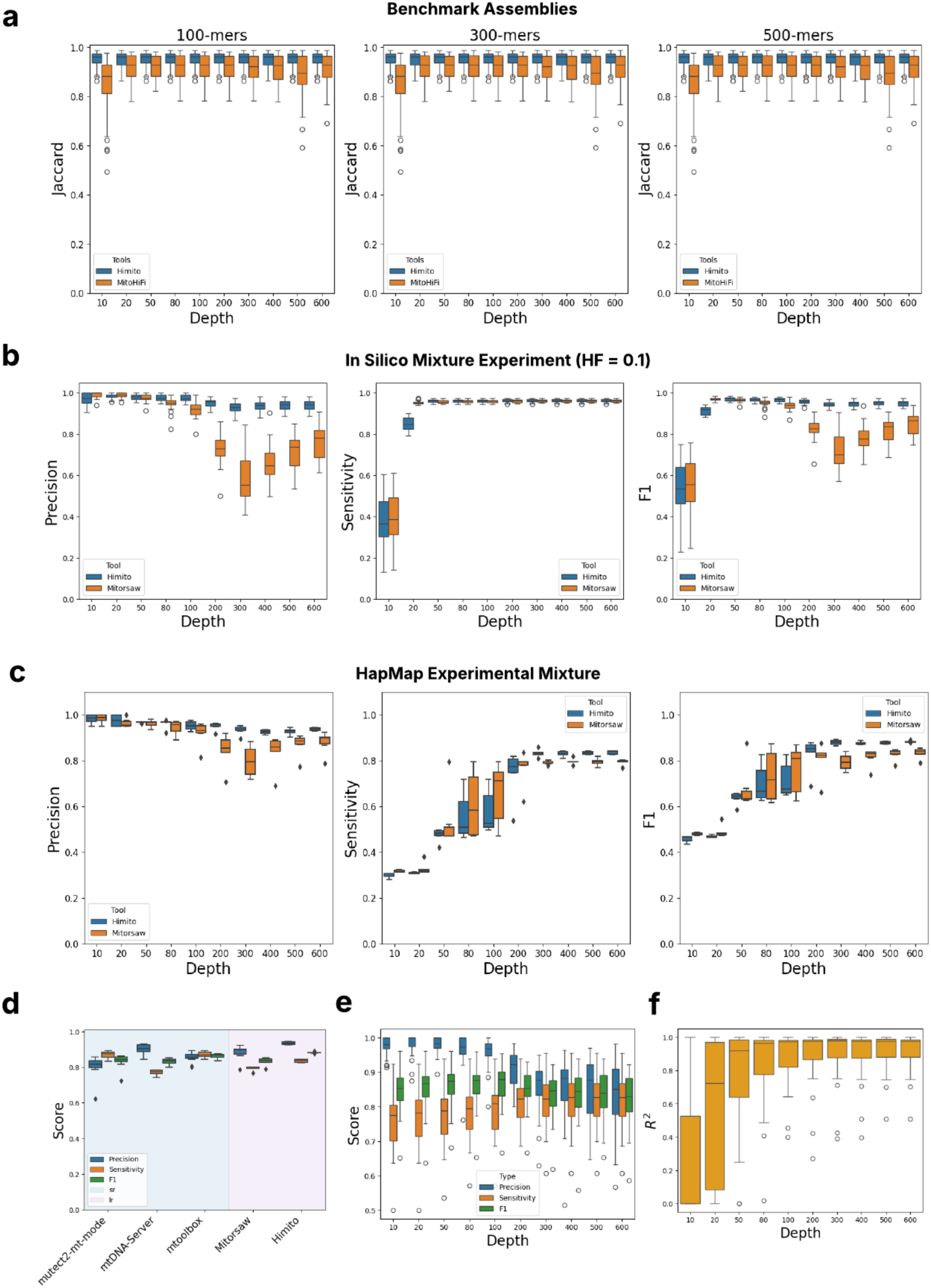
Benchmarking of *Himito* assemblies and variant calls. a, Assembly evaluation. *Himito* and *MitoHiFi* assemblies are compared with the high-quality HPRC reference assemblies. b, Callset evaluation at predefined heteroplasmic frequencies (HF = 10%) using *in silico* mixture experiment. *Himito* and *Mitorsaw* callsets were evaluated and compared in terms of Precision, Sensitivity and F1 score (full analysis on Supplementary Figure 4, 5). c, Long read callset evaluation using HapMap mixture data and the HapMap mitochondrial truth variant callset. d, Callset evaluation between short reads and long reads. HapMap short read and long read data are downsampled to 600x respectively, then called variants using *mtoolbox*, *GATK Mutect2-mitochondrial mode (mutect2-mt-mode)*, and *mtDNA-Server* for short reads (sr) and *Himito*, *Mitorsaw* for long reads (lr). e. Taking the HPRC full-coverage short read callset generated by GATK Mutect2-mitochondrial mode as the truth set, the Himito call performance is evaluated at different depths. f. The R^2^ values of allele frequency for true positive (TP) calls from srWGS and lrWGS were calculated.

We next evaluated the Call module’s performance. *Himito* `Call` outputs both homoplasmic and heteroplasmic variants in the standard VCF file with their heteroplasmic frequency (HF) recorded. Using the HPRC minigraph-cactus assembly-based VCF as the truth set, we assessed homoplasmic variants (HF > 95%) called by *Himito* using the standard VCF evaluation tool *RTG vcfeval*^34^. *Himito* achieved high precision and sensitivity (F1 ≈ 0.95; Supplementary Figure 3).

To assess heteroplasmic variant detection, we conducted an *in silico* mixture experiment simulating predefined heteroplasmic frequencies (Methods; Supplementary Figure 2). The assembly-based variants identified from the two donor samples were merged to create the truth set. We contrasted *Himito* performance with *Mitorsaw* (v0.2.0), a recently developed long-read mitochondrial variant caller. *Himito* outperformed *Mitorsaw* in variant calling sensitivity and overall F1 scores at lower heteroplasmic frequencies (1%-10% using default parameters) (Supplementary Figure 4). In addition, we found that *Mitorsaw’s* sensitivity to detect low-level heteroplasmic variants can be improved by adjusting the `minimum-maf` parameter, with the cost of decreased precision (Supplementary Figure 5). For a balanced comparison, we set the `minimum-maf` of Mitosaw and the `vaf-threshold` of *Himito* at the same value 0.01. As shown in Figure 2b, with a predefined frequency of 10%, both *Himito* and *Mitorsaw* show low F1 scores in low depth scenarios (10x), yet *Himito* outperforms Mitosaw when the depth is over 50x (complete results in Supplementary Figure 5).

We extended benchmarking of *Mitorsaw* and *Himito* to the HapMap mixture lrWGS dataset and its corresponding truth callset. This resource, developed by the SMaHT consortium^33^, was generated by experimentally mixing six HapMap cell lines^35^ to produce variants that span low (0.5–16.5% HF) and high (83.5–100% HF) heteroplasmy levels. The minigraph-cactus graph constructed from HPRC assemblies of these donors was used to derive the truth set.^36^ As shown in Figure 2c, *Himito* outperforms *Mitorsaw* in both precision and sensitivity using PacBio long-read dataset, as well as in overall F1 score, especially at high depth settings. In addition, we optimized the Himito parameters for ONT lrWGS analysis. We applied Himito to 5 ONT long-read dataset in the HapMap mixture. The results showed comparable performance with PacBio lrWGS data (Supplementary Figure 17). These results suggest that *Himito* shows better performance when calling mitochondrial variants compared to *Mitorsaw*.

To compare the variant calling performance between different data modalities, we downsampled the HapMap mixture short-read and long-read data to 600x and called variants using different short-read whole genome sequencing (srWGS) and lrWGS callers respectively. *Himito* shows higher precision and sensitivity among long-read callers (Figure 2d). Among short-read tools, *mtDNA-Server*^29^ demonstrated the highest precision, while *GATK-Mutect2* mitochondrial mode^28^ achieved the greatest sensitivity. Overall, long-read callers (*Himito*, Mitorsaw) excelled in precision, and in the case of Himito, matched or exceeded the sensitivity of the best short-read callers.

Finally, we compared *Himito* calls to the paired HPRC short-read callset generated by *GATK-Mutect2* mitochondrial mode. *Himito* achieved high F1 scores compared with short-read data as truth set (Figure 2e), and high concordance of heteroplasmic frequencies at depth exceeding 50x (Figure 2f). These results confirm good agreements between long-read and short-read results.

In addition, *Himito’s* methylation analysis aggregated 5mC signals along each major mitochondrial haplotype, enabling downstream methylation analyses. *Himito’s* methylation calling achieves high concordance with PbCpG-count mode (R^2^ > 0.99, Supplementary Figure 6).

### All of Us v8 PacBio Long Read Callset

We applied *Himito* to analyze the mitochondrial genetic variation in *All of Us* (AoU) v8 long-read dataset. The AoU v8 release incorporates 2,517 PacBio samples and 324 ONT samples. We assessed the sequencing depth across the mitochondrial chromosome for all PacBio samples and excluded those with an average depth below 10x. After filtering, 2,422 PacBio samples remained for downstream analysis with an average chrM depth of ∼80x (Figure 3a).

**Figure 3.**
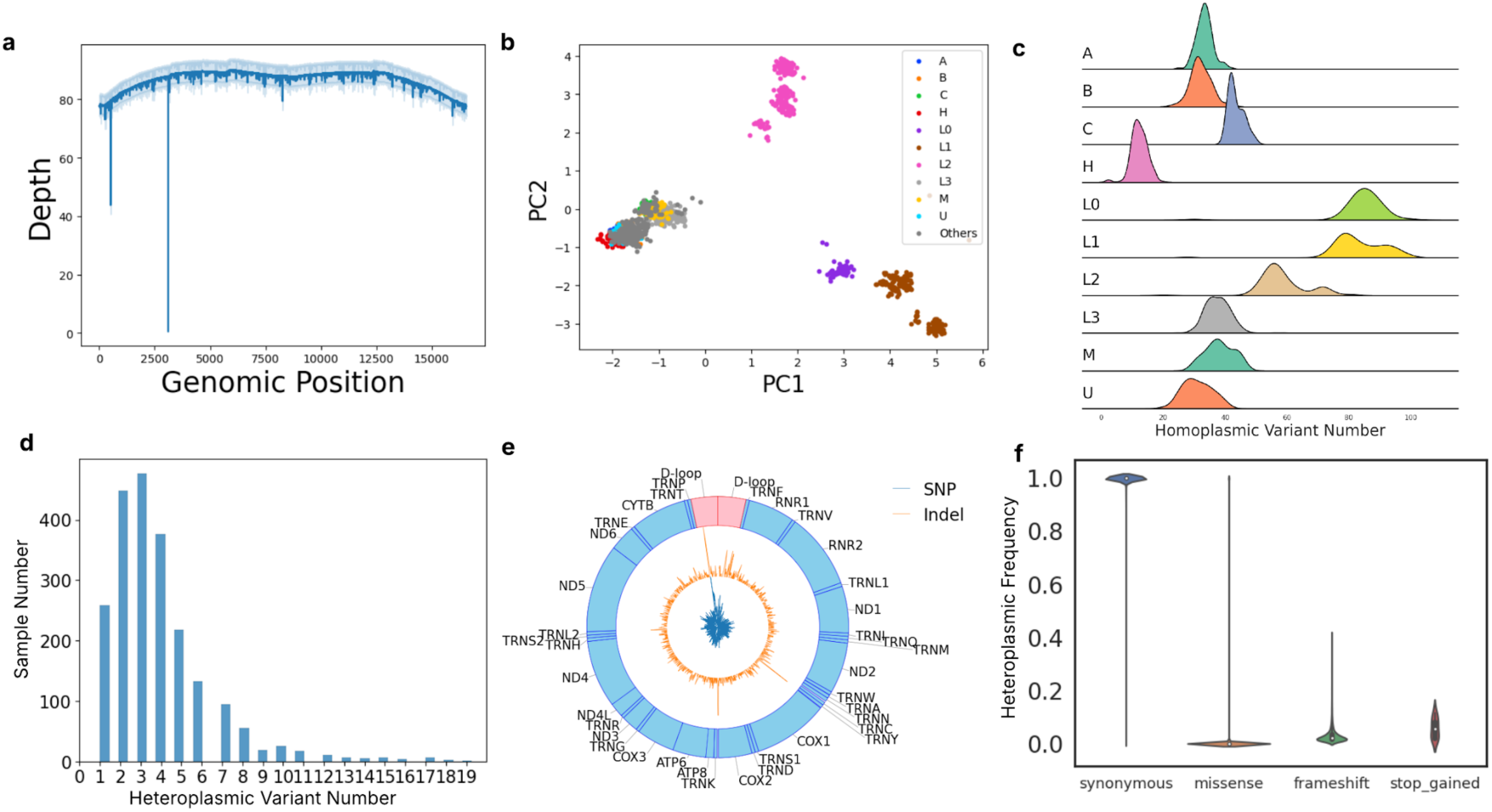
Genetic variants in AoU v8 PacBio samples. a. Coverage depth of mitochondrial genome (chrM) for the AoU v8 Pacbio samples. b. Principal Component Analysis (PCA) plot of mitochondrial genotypes across AoU v8 PacBio samples. c. Homoplasmic variant number distribution in different haplogroups. d. Heteroplasmic variant number distribution in AoU v8 PacBio samples. The mode of heteroplasmic variant number is 3. e. Variant density distribution, stratified by SNPs (blue) and Indels (orange) in the mitochondrial genome. f. Functional annotation of mitochondrial variants and their frequencies.

Mitochondrial variants were identified for each sample using *Himito* and subsequently merged with *bcftools merge*^37^. Haplogroups for each sample were determined using *haplogrep*^38^. Principal component analysis (PCA) was then conducted based on genotypes extracted from the *Himito* callset (Figure 3b). PCA revealed clear clustering of samples by their haplogroups, reflecting global population diversity. We then quantified the number of homoplasmic variants (HF > 95%) and heteroplasmic variants (HF between 1-95%) in each sample. The distribution of homoplasmic variant counts was multimodal, attributable to haplogroup diversity (Figure 3c), whereas heteroplasmic variant counts followed a unimodal distribution centered around three variants per sample (Figure 3d), consistent with the widespread presence of low-level heteroplasmies in human mtDNA.

We next examined the variant density (stratified by SNVs and indels) across the mitochondrial genome (Figure 3e). We observed the high concentration of genetic variations in the D-loop region in AoU callset, which agrees with the previous study of the presence of hypervariable regions in the mitochondrial genome^39^.

To characterize length heteroplasmies commonly arising from variable-length homopolymer tracts,^12^ we analyzed three known length heteroplasmic loci chrM:302-316 (ACCCCCCCTCCCCCG), chrM:567-574 (ACCCCCCA), and chrM:16174-16195 (CAAAACCCCCTCCCCA), and quantified the number of distinct haplotypes per sample. Joint haplotypes between each pair of these length heteroplasmies were analyzed (Supplementary Figure 13), and co-occurrence was further assessed at the read level using Fisher’s exact test (Supplementary Figures 14, 15). These analyses revealed that length-heteroplasmic variants often co-occur across distant genomic regions, suggesting shared mutational processes or linked replication dynamics.

We performed functional annotation of *Himito*-detected variants using the Variant Effect Predictor (VEP)^40^, classifying each variant by predicted consequence: synonymous, missense, frame shift, and stop-gain. Following annotation, we stratified variants based on their predicted consequences and analyzed their heteroplasmic frequencies (Figure 3f). Synonymous variants were most frequent, whereas missense variants, frameshift variants, and stop-gained variants tended to occur at lower frequencies, consistent with purifying selection acting on deleterious alleles.

We next intersected the *Himito* callset with the pathogenic variants reported in the MitoMap database^41^. Three pathogenic variants (m.1555A>G, m.11778G>A, and m.14484T>C) were identified among 2,422 individuals in the AoU v8 long-read cohort at heteroplasmic frequencies ranging from 10-100% (8 carriers total). These variants are associated with deafness, autism spectrum, atherosclerosis, Leber’s hereditary optic neuropathy (LHON), and progressive dystonia^41^. Examination of *All of Us* survey data confirmed that several high-frequency carriers reported related disabilities. Although low-frequency mitochondrial pathogenic variants may not cause overt disease, their identification in population-scale datasets may provide valuable opportunities for early detection and prevention.

### Pangenome of AoU v8 PacBio Assemblies

The development of third-generation sequencing technologies has enabled personalized genome construction and driven a paradigm shift from the linear reference genome to the pangenome models that integrate multiple genomes in a unified representation. Linear reference genomes have long served as the foundation of many bioinformatics pipelines, providing coordinate systems for read mapping, variant calling, and genome annotation. However, these linear reference genomes cannot capture the full breadth of human genetic diversity. In contrast, graphical pangenomes provide a comprehensive framework for representing sequence variation across populations.

Using *Himito*, we assembled individual mitochondrial genomes for 2,422 PacBio long-read samples from the AoU v8 long-read cohort and combined them to build a population-scale mitochondrial pangenome graph (Figure 4a). We showed the length difference between assembled and standard reference mitochondrial genomes (rCRS) in Figure 4b. Overall, the length differences are less than 50 bp, which indicates the absence of large indels in these primary mitochondrial genomes. The global topology of the population-scale pangenome graph is shown in Figure 4c. We constructed a binary matrix representing the presence or absence of each graph vertex within each individual genome. Hierarchical clustering of this matrix grouped samples by haplogroup (Supplementary Figure 7), confirming that the graph structure encodes haplogroup-specific mitochondrial diversity.

**Figure 4.**
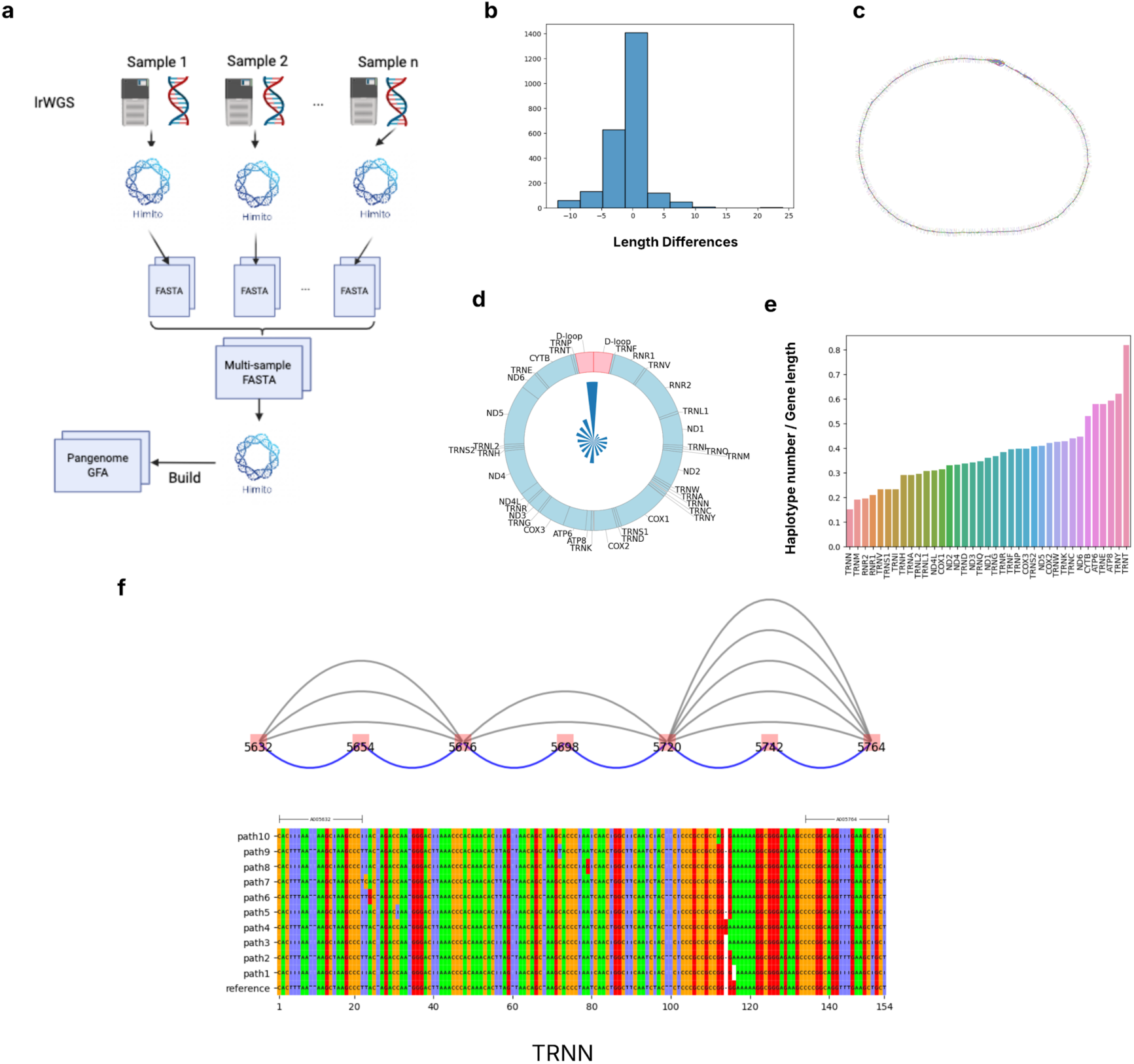
Pangenome for 2422 mitochondrial genome assemblies constructed from AoU v8 PacBio samples. **a.** Pangenome graph construction workflow. **b.** Length difference between *Himito* assemblies and the standard human mitochondrial reference genome (rCRS). **c.** Global structure of the mitochondrial pangenome constructed from individual assemblies. **d.** The number of distinct haplotypes within each 1 kb window in the mitochondrial pangenome. **e.** Sequence diversity scores for each mitochondrial gene, measured as the number of unique haplotypes normalized by the gene length. f. The local graph structure in the TRNN gene. Multiple sequence alignment was performed among all the unique haplotypes in the gene.

Local subgraphs illustrate regional sequence variation within specific genomic intervals. Leveraging the anchor-based sequence coordinate system, we identified subgraphs and haplotypes within predefined regions (Methods). For each non-overlapping 1 kb window, we enumerated all unique haplotypes across the mitochondrial genome. The D-loop region exhibited a significantly higher number of haplotypes than any other region in the mtDNA. In contrast, MT-RNR1 and MT-RNR2 gene regions, which encode the ribosomal RNAs, are relatively conserved across the 2,422 individuals. For each gene region annotated on the standard reference genome, we computed a sequence diversity score, defined as the number of distinct haplotypes for each gene normalized by the gene length (Figure 4e).

As an example, we visualized the local graph structure for the TRNN gene, encoding mitochondrial tRNA gene encoding transfer RNA for asparagine (Asn) (Figure 4f). Multiple sequence alignment among haplotypes of this gene revealed base-level polymorphisms within this region, illustrating how the pangenome graph provides an intuitive and comprehensive view of conserved and variable features across the population. Together, these analyses demonstrate that the mitochondrial pangenome graph captures both global and local sequence diversity across individuals.

### Methylation Signals in the Mitochondrial Genome

Among the *All of Us* v8 PacBio long-read samples, 1,490 samples contained methylation data. We investigated the methylation patterns in the 1,490 PacBio long-read samples and 324 ONT samples from the AoU v8 cohort. Reads aligned to chrM were stringently filtered by their alignment signatures and methylation levels. Reads with over 20% of their CpG sites with a methylation likelihood greater than 0.5 were stringently filtered out as potentially derived from NUMTs (Figure 1a, Supplementary Methods).

After filtering, we calculated the modification rate of each CpG site using *Himito* `methyl`. Despite stringent filtering, methylated CpGs were consistently observed across the mitochondrial genome in both PacBio and ONT samples. However, notable technical variation existed between PacBio and ONT data (Supplementary Figure 8, 41 paired PacBio and ONT samples in v8). To account for this, we applied distinct modification rate thresholds (0.3 for PacBio and 0.1 for ONT) determined from the overall modification rate distributions (Supplementary Figure 9).

We next examined the distribution of hypermethylated CpG sites across individuals (Supplementary Figure 10). Several CpG sites appeared recurrently hypermethylated in multiple samples. Overall, these findings provide evidence supporting the presence of 5mC methylation modification in the mitochondrial genome, while also highlighting technical variability across sequencing technologies.

### Age-Related Observations

The AoU v8 long-read cohort represents a large and diverse population with rich phenotypic data, including participant age, which ranges from 20 to 90 (Supplementary Figure 11). A recent study, using srWGS data from AoU, showed that an increase with age in low heteroplasmy mtDNA SNVs was due to a reduction in blood cell clonal diversity after middle age that in turn increases the sensitivity of bulk sequencing data for detecting the mtDNA variants of the dominant clones^42^. We found a similar age-associated accumulation of low-frequency SNVs (1-10% heteroplasmic frequency) in the lrWGS data (Figure 5a). Our results also validate that the increase in low heteroplasmy SNVs with age is dominated by G>A and T>C transitions, consistent with these variants being introduced by a replication-linked mtDNA mutational process^42^ (Figure 5b; Supplementary Figure 12). We also observed a decrease in the number of distinct haplotypes with age in the chrM:302-316 repeat expansion region (Supplementary Figure 15).

**Figure 5.**
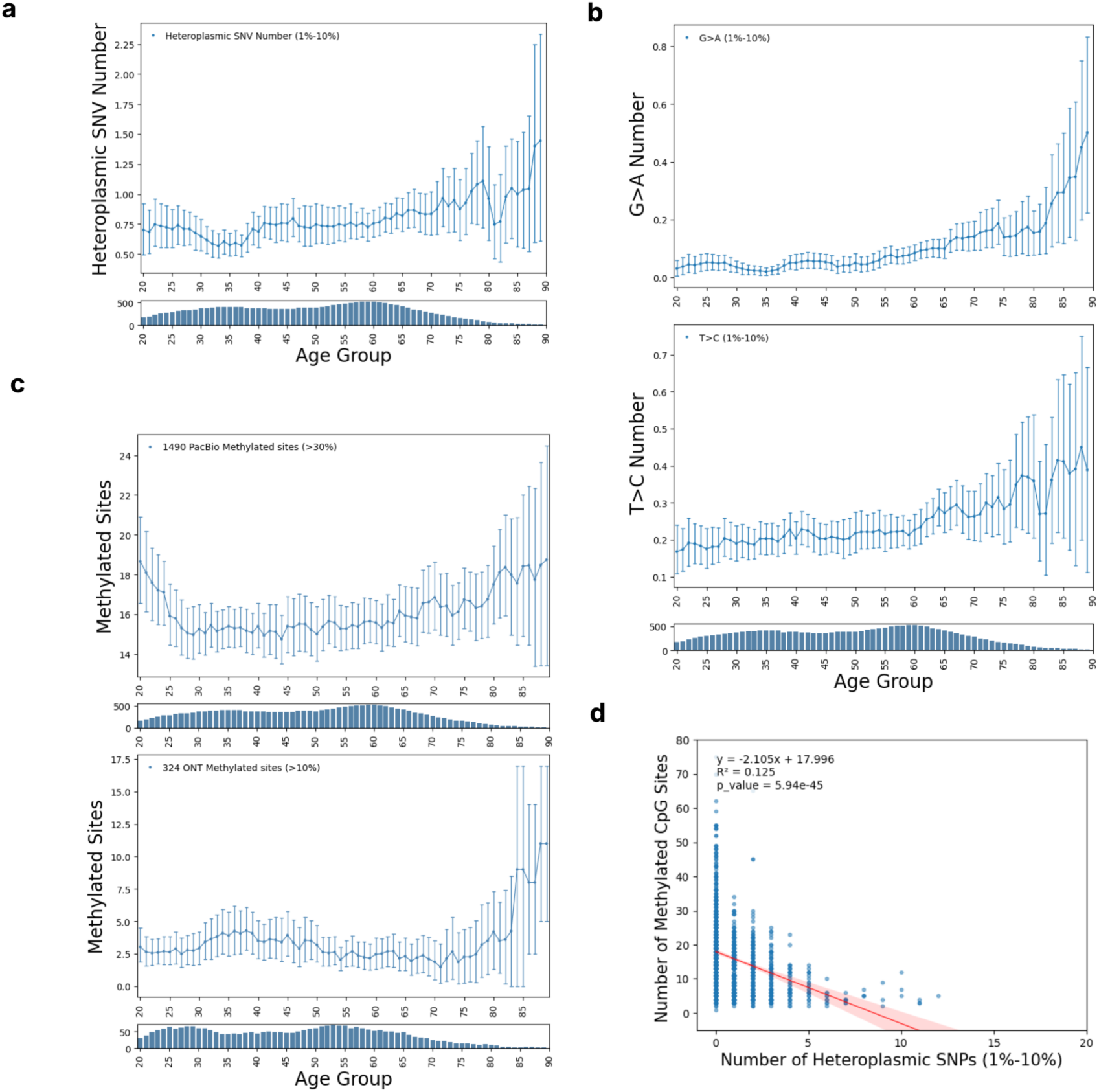
Age-related observations. a. The number of low-frequent heteroplasmic SNVs (1-10%) accumulated with age (sliding window size of 10). The number of individuals in each bin is shown in the bottom plot. b. A subset of low-frequent heteroplasmic SNVs, G to A and T to C shows the age-accumulating patterns.The full analysis shows in Supplementary Figure 12. c. The number of hyper-methylated CpG sites (mod rate > 0.3) in the 1490 PacBio samples and in the 324 ONT samples (mod rate > 0.1) stratified by age groups. d. Negative correlation between the number of low-frequent heteroplasmic SNVs (1-10%) and the number of methylated CpG sites (Mod score > 30%) in PacBio samples (P value = 5.94e^-45^).

Intriguingly, analysis of both PacBio and ONT methylation data revealed an increase in the number of hypermethylated CpG sites with age (Figure 5c). Moreover, a significant negative correlation exists between the counts of low-frequency SNVs and methylated sites per sample, including when correcting for age (Figure 5d, p-value < 1e-44 after age correction, age p-value < 1e-02). A prior study showed that *de novo* mtDNA methylation prevents stress-induced mtDNA damage during the critical peri-implantation period of embryogenesis^43^, which provides a possible explanation of the antagonistic relation between methylation and low-frequency SNVs in our data. Himito’s ability to jointly analyze variation and methylation levels could enable future explanation of the role of mtDNA methylation in mitochondrial biology.

## Discussion

In this study, we present *Himito*, a graph-based toolkit designed for mitochondrial genomic and epigenomic analysis using long reads. *Himito* addresses several key challenges in mitochondrial genomics, including resolving the circular structure of mtDNA, removing the NUMTs-derived reads, identifying low-level heteroplasmies, and integrating 5mC methylation signals. *Himito* integrates both genomic and epigenomic features in a single graph representation, and excludes reads derived from NUMTs that have been misaligned to improve the accuracy of the downstream genome assembly, variant calling, and methylation analysis. The *Himito* graph reveals the circular nature of mtDNA, which facilitates visualization of sequence diversities. The haplotypes, sequence variants, and methylation signals can be extracted from the graph in standard file formats, which are compatible with standard bioinformatics tools. *Himito* is written in Rust, which is easy to install and integrate into existing bioinformatics pipelines.

Compared with existing tools, *Himito* delivers more accurate mitochondrial genome assemblies, improved heteroplasmic variant detection, and integrated methylation analysis. Benchmarking against the state-of-the-art methods using publicly available HPRC and HapMap mixture lrWGS dataset demonstrated that *Himito* yields higher assembly accuracy than *MitoHiFi*^31^ and higher sensitivity for low-frequency heteroplasmic variants than *Mitorsaw*. Together, these results establish *Himito* as a powerful and versatile toolkit for comprehensive long-read mitochondrial genome analysis.

Application of *Himito* to the *All of Us* long-read dataset enabled population-scale investigation of mtDNA structure, variation, and methylation. Analysis of the AoU mitochondrial PacBio long-read callset elucidates the haplogroup composition within the AoU cohort and identifies pathogenic variants in the AoU v8 samples. We then assembled the major haplotype for each sample, constructed a population-level pangenome graph to represent 2,422 genomes in the AoU v8 cohort. We investigated the conserved and highly diverse regions in the mitochondrial genome. The sequence graph provides an intuitive visualization of sequence diversity in a specific genomic region and enables a better understanding of the genomic structure within the mitochondrial genomes. Our analysis of mtDNA methylation further suggests that, despite its low level, 5mC methylation exists in the mitochondrial genome. We investigated the age-related genomic and epigenomic patterns of the mitochondrial genome and validated an increase in low-heteroplasmy SNVs with age present in srWGS data that was shown to be the result of reduced blood cell clonal diversity with age,^42^ and further identified a similar trend in the number of hypermethylated CpG sites. We observed a negative correlation between the number of low-frequency SNVs and methylated CpGs, suggesting a potential interplay between these processes. These findings require further investigation into the underpinning mechanisms and their functional roles. Himito facilitates the integrative analysis of mitochondrial genomes, demonstrating the effectiveness of combining variation and methylation analyses through lrWGS.

Despite these strengths, there are several limitations to consider. First, *Himito* is capable of detecting low-frequency heteroplasmies with sufficient coverage; however, caution should be taken when distinguishing low-level heteroplasmic variants from sequencing errors. Orthogonal validation using multiple technologies, such as integrating short-read and long-read data, is helpful for confirming the presence of low-heteroplasmic variants within a single sample. For improved mitochondrial genome analysis with *Himito*, the raw variant callset from AoU v8 requires the implementation of more advanced filtering techniques before downstream association analysis. In addition, *Himito’s* variant calling performance on lower coverage or noisy datasets may be reduced. *Himito* only incorporates 5mC modification in the analysis. The accuracy of epigenetic modification detection is dependent on the sequencing platform, the methylation caller, and the sample preparation. More comprehensive studies on the technical variations in methylation analysis are needed to be performed. Future work will aim to address these challenges by incorporating more sophisticated variant filtering strategies, supporting hybrid data integration, and expanding compatibility with additional epigenetic modification types.

As long-read sequencing becomes increasingly cost-effective, population-scale datasets will continue to expand. *Himito* provides a scalable foundation for exploring mitochondrial disease mechanisms, age-related processes, and tissue-specific variation across large cohorts such as the *All of Us* and SMaHT^44^. Using appropriate reference genomes and long-read sequencing data as input, *Himito* can be applied beyond human mitochondrial genomes to those of other species. For example, *Himito* can build a cross-species pangenome model designed to enhance the understanding of mitochondrial genome evolution across species, utilizing data obtained from consortium efforts such as the Darwin Tree of Life Project^45^.

In summary, *Himito* expands the toolkit available for mitochondrial genome analysis by introducing a robust, accurate, and methylation-aware graph-based framework tailored to the features of mtDNA and long-read sequencing. This unified approach enables integrated investigation of mitochondrial genetic and epigenetic diversity and offers a foundation for future studies of disease, aging, and evolution.

## Data Availability

*Himito* is publicly available at https://github.com/broadinstitute/Himito. The HPRC data is available in https://anvil.terra.bio/#workspaces/anvil-datastorage/AnVIL_HPRC. The AoU v8 release can be accessed via research workbench https://www.researchallofus.org/data-tools/workbench/. The AoU v8 lrWGS analysis is performed in Research Workbench Workspace: https://workbench.researchallofus.org/workspaces/aou-rw-2cf44b8f/aoulrmitochondrialgenomeanalysis/data. The HapMap dataset can be found in the SMaHT data portal at https://data.smaht.org/.

## Acknowledgements

Special thanks to **Vamsi Mootha** and **Sarah Calvo** at Broad Institute for their helpful discussion and suggestions to the development and evaluation of *Himito*. Many thanks to **Na Cai** at ETH Zurich for her suggestions testing the heteroplasmic variants co-occurrence. Many thanks to **Kristin Ardlie, Tim Coorens** from Broad Institute for their support for the SMaHT Pilot and Feasibility award application. We thank **James Emery, Ryan Lorig-Roach, Fabio Cunial, Lucas van Dijk, Shadi Zaheri** at Long Read lab, Broad Institute for their valuable discussions during the development of *Himito*. Thanks to **Data Science Platform** for the use of computational infrastructure provided by Terra. Thanks to the **SMaHT Pilot and Feasibility Award** for supporting this project.

## Author Contributions

H.S., Y.H., and K.V.G. conceived the project and designed the algorithm. H.S. developed the toolkit and performed the benchmarking and AoU lrWGS analysis. T.D. contributed to the interpretation of AoU lrWGS callset discovery and age-related pattern. N.K., E.C., S.C.J. contributed to HapMap mixture dataset development and validation. N.K., E.C. contributed to the srWGS benchmarking using HapMap dataset. D.B. contributed to the variant filtering algorithm. K.V.G. and Y.H. supervised the project.

## Methods

### Algorithms

#### *Himito* includes five key modules for comprehensive mitochondrial genome analysis

`Filter` (Figure 1a): The Filter module aims to remove reads from NUMTs to ensure the accuracy of downstream analysis. It classifies reads using both alignment signatures (SA tags in BAM files) and read-level methylation patterns, identifying and separating NUMT-derived reads from mtDNA-derived reads.

`Build` (Figure 1b): The Build module focuses on constructing a graphical genome model for mtDNA. It selects unique k-mers as anchors in the reference genome, maps these to long reads, extracts sequences between anchor pairs, consolidates identical sequences into graph vertices, and automatically forms a circular graph representing the mtDNA.

`Asm` (Figure 1c): The Asm module is designed to assemble the primary mitochondrial haplotype from the constructed graph. It reconstructs the main haplotype sequence by traversing the graph and selecting the most supported paths, then outputs these sequences in FASTA format.

`Call` (Figure 1d): The Call module detects both homoplasmic and heteroplasmic variants from the graph. It enumerates paths between anchor pairs, aligns these paths to the reference segments, records alignment differences as cigar strings, maps variants to reference coordinates, and outputs them in VCF format, which includes read counts, depths, and heteroplasmic frequencies information, while also employing a permutation test to filter false positives.

`Methyl` (Figure 1e): This module analyzes DNA methylation at both read and haplotype levels. It identifies 5mC modifications at CpG sites using MM and ML tags, annotates methylation likelihoods to graph vertices, aggregates methylation signals, estimates methylation rates for CpGs on the major haplotype, and outputs these rates in BED format.

Please see the supplementary file for detailed description.

### Permutation Test for Variant Filtering

One major challenge for heteroplasmic variant calling is to distinguish the low level heteroplasmic variants with sequencing artifacts. We adopted a novel strategy to exclude false positive calls based on the co-occurrence patterns of variants on long reads. The sequencing artifact, especially the extensively observed short indels in both the PacBio and ONT long read sequences, randomly occurs among all of the reads. With the assumption that the true variants co-occur with other genuine variants in the same haplotype, we performed the permutation test to filter out variants that are likely to be sequence artifacts. A presence-absence matrix was initially built to represent each variant’s occurrence across all reads. We calculated the sum of each pair-wised Jaccard similarity of variants as the statistics. By randomizing the variants occurrence in the reads, we calculated the null distribution by shuffling the occurrence of the variants in all reads and computing the sum of all pair-wised Jaccard similarity. We then computed the empirical p-values for each observation to exclude the variants that agree with the null hypothesis. This approach effectively excluded many false positive calls in the raw graph and improved the precision of the *Himito* heteroplasmic variant calling.

### In silico Mixture Experiment to Evaluate Heteroplasmic Calling Performance

To evaluate the precision and recall of variant calling at a defined heteroplasmic frequency, an *in silico* mixture experiment was conducted (Supplementary Figure 2). This experiment utilized the sequencing reads aligned to the mitochondrial chromosome (chrM) from two distinct donor samples, which involves distinct sets of true variants. For each donor, the aligned BAM file was downsampled to a specified coverage depth. Subsequently, the downsampled BAM files from the two donors were merged at a predefined mixing ratio, such as 0.1 and 0.9, to simulate a sample with a known heteroplasmic variant frequency. Variant calling was then performed on the resulting mixed BAM file. The truth set for variant evaluation was established as the union of all assembly-based variants identified in the two donor samples. Finally, the performance of the variant caller on the mixed sample was assessed by comparing the called variants against the truth set using the *RTG vcfeval* tool, allowing for the quantification of precision and recall at the simulated heteroplasmic frequency.

The workflow is in https://github.com/broadinstitute/Himito/blob/main/wdl/MixSampleExperiment.wdl

HapMap Mixture Experiment and Mitochondrial Truth Callset Generation

The SMaHT^33^ network created an in vitro-constructed cell line mixture using six well-characterized cell lines from the International HapMap Project^35^, composed of 83.5% HG005, 10% HG02622, 2% each of HG002, HG02257, and HG02486, and 0.5% HG00438. Genome Characterization Centers at Baylor College of Medicine, the Broad Institute, the University of Washington and Seattle Children’s Research Institute, and Washington University in St. Louis generated high-depth PacBio long-read whole-genome sequencing (WGS) data for this mixture. The HapMap mixture somatic variant truth set^36^, including mitochondrial variants, was constructed using a minigraph–cactus pangenome graph built from de novo assemblies produced by the Human Pangenome Reference Consortium^46^. The aligned BAM files and the corresponding truth set were obtained from the SMaHT data portal.

For each PacBio long read whole genome sequencing sample in the HapMap dataset, we first subset the whole genome BAM file to reads aligned to chrM. We downsampled the BAMs to 10x, 20x, 50x, 80x, 100x, 200x, 300x, 400x, 500x, 600x, then called *Himito* and *Mitorsaw* at each coverage level respectively to generate the mitochondrial callset. We used RTG vcfeval to evaluate the performance of *Himito* and *Mitorsaw* respectively. We further benchmark ONT long read whole genome sequencing samples in the HapMap dataset using the same strategy. The analysis workflow is in https://github.com/broadinstitute/Himito/blob/main/wdl/HapMap_downsample_experime <u>nt.wdl</u>

### Population-level Pangenome Construction

Mitochondrial genomes were assembled by *Himito* for each sample in AoU v8 release individually using PacBio long reads. These assemblies were then combined and used as input for the *Himito* `Build` function to construct a population-level genome graph. This graph was further processed and circularized using customized scripts. For specific genomic regions of interest, the number of distinct haplotypes represented in the graph were extracted and analyzed to assess regional diversity. Anchors served as a sequence-based coordinate system. We associated each graph vertex to a reference interval utilizing the bounding anchor sequences and their reference coordinates. For each genomic region, such as genes, we found all the graph vertices overlapping with the interval and reconstructed the subgraph in that genomic region. We then traversed the subgraph and extracted all the haplotypes in that region, without any coverage-based trimming.

The distinct haplotypes within each genomic region were recorded in the standard FASTA files. Mitochondrial haplotype differences were analyzed using `ClustalW`^47^ for multiple-sequence alignment (MSA) and MSAplot (https://github.com/mourisl/MSAplot) for visualization. Haplotype diversity was quantified by counting distinct haplotypes in 1kb windows across the genome and visualized with `pyCirclize`^48^. Gene intervals from the rCRS GFF3 file (NCBI) were used to extract subgraphs from the AoU v8 pangenome. Distinct haplotype sequences were identified by graph traversal within these subgraphs. Genetic diversity per gene was calculated as the number of distinct haplotypes normalized by gene length, providing a measure of gene-specific diversity.

