## Supplementary file for "*Himito*: a Graph-based Toolkit for Mitochondrial Genome Analysis using Long Reads"

### Himito Supplementary

Table of contents for supplementary Materials


Supplementary Methods 2
- Detailed Algorithm Description 2
  - Distinguishing Mitochondrial DNA (mtDNA) from Nuclear Mitochondrial Sequences (NUMTs) 2
  - Constructing Graphical Genome from mtDNA-derived Reads 3
  - Extracting Primary Haplotype from the Himito Graph 4
  - Calling Homoplasmic and Heteroplasmic Variants from the Himito Graph 4
  - Variant Filtering Strategies 5
  - Annotating Methylation Signals to the Himito Graph and Signal Aggregation along the Major Haplotype 6
- Benchmarking Methods 7
  - Assembly evaluation using HPRC truth assemblies 7
  - In silico Mixture Experiment for Heteroplasmic Variant Calling Evaluation 9
  - Benchmarking Long-read Callers using HapMap Mixture Dataset 13
  - Benchmarking against short read sequence data 15
    - Evaluation between short-read and long-read data using HapMap Mixture 15
    - Concordance between short-read and long-read callset 16
  - Methylation Calling Evaluation Methods 18
- AoU v8 Callset Analysis 19
  - Sample Depth Analysis 19
  - Variant Calling and Data Processing for AoU v8 PacBio Samples 20
  - Haplogroup Classification 21
  - Functional Annotation using VEP 22
  - Pathogenic Variants analysis 23
  - Length Heteroplasmic Variants Analysis 23
- AoU v8 Pangenome Construction and Analysis 23
  - AoU v8 Pangenome Construction 23
  - Haplotype Number determination 26
  - Sequence Diversity Score Calculation 27
- Detailed Explanation of Methylation Data Processing 27
- Methylation Analysis in the All of Us (AoU) v8 Long-read Release Using PacBio and ONT 29
- Age-related Analysis 30

Supplementary Figures 31
- Supplementary Figure 1 Workflow for Himito. 32
- Supplementary Figure 2 In silico mixture experiment for heteroplasmic calling evaluation. 33
- Supplementary Figure 3 Homoplasmic calling evaluation. 34
- Supplementary Figure 4 Variant calling benchmark against Mitorsaw at default parameters

(Mitorsaw minimal-MAF = 0.1, Himito vaf-threshold = 0.01). 35
Supplementary Figure 5 Mitorsaw and Himito comparison with the same detection limits (Mitorsaw minimum-maf = 0.01, Himito vaf-threshold = 0.01). 36
Supplementary Figure 8, PCA plot for CpG modification rate in 41 paired PacBio and ONT samples in AoU v8 cohort. 39
Supplementary Figure 9, The modification rate distribution. 40
Supplementary Figure 10, Recurrent hypermethylated CpG sites across PacBio (modification rate threshold = 0.3) and ONT samples (modification rate threshold = 0.1). 40
Supplementary Figure 12. The number of different variant types in Age groups (sliding window of 10). 43
Supplementary Figure 13. Number of length heteroplasmic variants per individual and the joint alleles distribution. 43
Supplementary Figure 14. Fisher's Exact test for the association for length heteroplasmic alleles across 229985 reads. 45
Supplementary Figure 15 Co-occurrence of length heteroplasmies. 45
Supplementary Figure 16 Haplotype Number of chrM:302-316 in Age groups (sliding window of 10). 46
Supplementary Figure 17 HapMap mixture benchmarking using different long read sequencing technology (PacBio, ONT). 46

### Supplementary Methods

#### Detailed Algorithm Description

##### Distinguishing Mitochondrial DNA (mtDNA) from Nuclear Mitochondrial Sequences (NUMTs)

The initial step in the Himito pipeline involves distinguishing the mtDNA-derived reads from the NUMTs-derived reads. Himito extracts sequencing reads that align to the mitochondrial chromosome (chrM). Following this step, Himito identifies reads that possess a supplementary alignment to any autosomal chromosome. These reads are flagged as originating from NUMTs. This filtering process is implemented by systematically scanning the Supplementary Alignment (SA) tag present in the standard BAM (Binary Alignment Map) file format. By examining the SA tag for evidence of supplementary alignments to autosomal loci, Himito effectively distinguishes true mitochondrial reads from those NUMTs-derived reads that may confound the analysis due to their nuclear origin.

Secondly, long-read sequencing technologies like PacBio and ONT generate both genetic sequence and methylation data. Existing research suggests that mitochondrial DNA has lower 5mC methylation compared to nuclear DNA. Our analysis of per-site methylation likelihood shows distinct distributions between nuclear and mtDNA reads (Figure 1a, PacBio HiFi reads). By calculating the percentage of methylated sites per read, we observed that reads containing nuclear DNA exhibit a higher percentage of methylated sites. Consequently, we exclude reads exceeding a maximum methylation percentage (default 0.2). This methylation threshold can be adjusted according to the observed methylation distribution in the long read data.

Finally, the identified NUMT-derived reads and mtDNA-derived reads are saved into separate BAM files in the output. To ensure the accuracy of downstream mitochondrial DNA analysis, the NUMT-derived reads are subsequently excluded from all further processing steps within the Himito pipeline.

#### Constructing Graphical Genome from mtDNA-derived Reads

An anchor-based graphical genome is constructed through the integration of a standard reference genome with long reads identified as mitochondrial DNA (mtDNA) derived in each sample. Unique k-mers within the standard mtDNA reference genome are selected as anchor sequences.The resolution of the sequence graph is controlled by the parameter k, which typically falls between 15 and 31, and dictates the length of anchor. Adjacent or overlapping anchor sequences are eliminated. These anchor sequences are mapped onto each of the long reads, and their respective mapping positions are recorded. Sequences located between pairs of anchor sequences are extracted, and identical sequences are consolidated into a single vertex within the graph. The resulting graph is stored as a standard GFA file, wherein 'S' and 'L' entities denote graph vertices and graph connectivities, respectively. Reference coordinates are annotated to the anchor sequences. Read names that support each path are also annotated to the graph. All annotations are documented in JSON format. In essence, the anchor sequences facilitate the organization of mtDNA-derived long reads into a sequence graph. This graph automatically forms a circular structure, which reflects the circular nature of mtDNAs (Figure 1b). A reference path traverses anchor pairs, enabling the mapping of genomic features, such as genes or exons, onto the graph. The local topology of the graph represents sequence diversity within a specific genomic region.

#### Extracting Primary Haplotype from the Himito Graph

The Himito sequence graph is a comprehensive representation of mitochondrial DNAs, integrating both the nucleotide sequence and associated annotations for every path within the graph. These annotations, recorded in the computer-readable JSON format, provide genomic information, including the standard reference coordinates for each anchor sequence, the specific set of long reads that support each consolidated path, and a detailed map of the connections between the different vertices within the graph. The primary haplotype sequence for each sample can be reconstructed by traversing the graph following the paths with the highest support from the input reads. (Figure 1c). These primary haplotype sequences are then exported in the standard FASTA file format, ensuring compatibility with existing bioinformatics tools and workflows. All primary haplotypes reconstructed from the `Himito Asm` initiate at the same point within the mitochondrial genome. This common starting location corresponds to the first anchor, defined as the first unique 21-mer identified within the revised Cambridge Reference Sequence (rCRS) reference genome.

#### Calling Homoplasmic and Heteroplasmic Variants from the Himito Graph

Homoplasmic and heteroplasmic variants are naturally represented in the alternative paths in the Himito graph. We enumerated all the paths between a pair of anchors and aligned these paths to the mtDNA reference sequence between anchor pairs. We excluded paths supported by fewer than a minimal count of reads (default: 2), as they were likely due to sequencing errors. We performed affine-gap alignment[36] between the reference and alternative segments and recorded the alignment as a cigar string[37] in each path, where “Ms” represent matches, “Xs” represent mismatches, “Ds” represent deletions and “Is” represent insertions. Compressed edits in the cigar strings, including mismatches, insertions and deletions, are mapped back to the standard reference coordinates and reformatted into a standard VCF file. Each record is annotated with the read counts supporting the alternative alleles (RC), read depths over a specific position (DP), and heteroplasmic frequency (HF). Along with the standard VCF file, the output includes an annotated graph GFA file that incorporates cigar strings. The presence or absence of each variant on every read is also provided as a binary matrix.

#### Variant Filtering Strategies

Distinguishing genuine low-level heteroplasmic variants from sequencing artifacts is challenging in heteroplasmic variant calling. To address this challenge, we developed a novel strategy that leverages the co-occurrence patterns of variants observed on long-read sequencing data. A key characteristic of sequencing artifacts, particularly the short indels frequently appearing in both PacBio and Oxford Nanopore Technologies (ONT) long reads, is their random distribution across all reads. Our approach operates under the assumption that authentic variants tend to co-occur with other genuine variants within the same haplotype. We employed a permutation test to filter out variants that were likely sequencing artifacts. This test involved several steps. First, we generated a binary matrix representing the presence or absence of each variant within each individual read. Next, we calculated the pairwise Jaccard similarity for every pair of variants, summing these similarities to derive a single statistic. This sum served as our observed value against which we would compare the results of our permutation test.

To establish a null distribution, we simulated the random occurrence of variants across the reads. This was achieved by shuffling the presence/absence data for variants in all reads, thereby randomizing their co-occurrence patterns. Following each shuffle, we recalculated the sum of pairwise Jaccard similarities, generating a set of values that represented the expected distribution if variants occurred independently of each other. By repeating this shuffling process many times, we obtained a robust null distribution.

With the null distribution established, we then computed empirical p-values for each observed statistic. These p-values allowed us to assess the likelihood of observing the actual co-occurrence patterns under the null hypothesis of random occurrence. Variants that exhibited co-occurrence patterns consistent with the null hypothesis (i.e., those with high p-values) were deemed likely to be sequencing artifacts and were subsequently excluded from the analysis. This process effectively eliminated numerous false positive calls that were present in the initial, unfiltered data.

The implementation of this permutation-based filtering strategy significantly improved the precision of Himito heteroplasmic variant calling. By focusing on co-occurrence patterns and statistically testing against a null distribution, we were able to refine our variant calls and substantially reduce the number of false positives, leading to a more accurate representation of true heteroplasmic variation.

#### Annotating Methylation Signals to the Himito Graph and Signal Aggregation along the Major Haplotype

Himito's methyl function provides a read-level and haplotype-level analysis of DNA methylation using long-read sequencing data stored in BAM files. By examining the MM and ML tags within each alignment record in the BAM files, Himito identifies individual 5mC modification at CpG dinucleotides. For each identified CpG site, Himito annotates the methylation likelihood, representing the probability of that specific cytosine being methylated to the corresponding vertex in the sequence graph. The information is stored in a JSON format in the standard GFA file, allowing for easy integration with other graph-based analyses. Following the initial mapping and likelihood assignment, Himito performs an aggregation of these methylation signals at each vertex of the graph. This aggregation step pools the information from multiple reads spanning the same CpG site, enabling the methylation rate estimation. Subsequently, Himito calculates the methylation rate of CpGs on the major haplotype in the standard BED file. For each CpG site present along the primary haplotype, Himito determines the number of methylated CpGs by a predefined threshold (default 0.5) and then calculates a modification rate, representing the proportion of reads showing methylation at that specific location. Finally, these calculated methylation rates are outputted in the standard used BED file format. The resulting BED files typically contain information such as the chromosome, start and end coordinates of the CpG sites, and the calculated methylation rate, providing a detailed and readily accessible summary of the methylation signals on the mitochondrial genome. This standard format allows for downstream integration and visualization of the methylation data with various genome browsers and downstream analysis pipelines. In addition to the BED file, Himito also outputs a *CSV* file containing the information of read-level methylation signals in the standard reference coordinates.

### Benchmarking Methods

#### Assembly evaluation using HPRC truth assemblies

This section details the methods used to benchmark the performance of Himito in generating mitochondrial assemblies, comparing it against MitoHiFi and a high-confidence truth set derived from the Human Pangenome Reference Consortium (HPRC) and HapMap mixture data. The workflow is at https://github.com/broadinstitute/Himito/blob/main/wdl/AssemblyBenchmark.wdl

The HPRC dataset, which includes high-quality assemblies for 47 individuals from diverse populations in the year-1 release, was utilized as a benchmark dataset. Specifically, the mitochondrial chromosome (chrM) contigs from the HPRC assemblies were extracted and employed as the truth set for assembly evaluation. These truth assemblies were identified by initially aligning whole-genome assemblies to a standard reference genome (GRCh38), followed by subsetting to the primary contigs that aligned to the mitochondrial chromosome (chrM). A standard reference genome (rCRS) for the mitochondrial chromosome was also used in the assembly processes for both Himito and MitoHiFi.

MitoHiFi assemblies were generated using the following sequential commands:

Step 1:

```
samtools view -bhX {whole-genome-bam} {whole-genome-bai} {locus} >
{prefix}.bam
```

This step extracts reads mapping to the mitochondrial locus from the input lrWGS BAM file.

Step 2:

```
samtools index {prefix}.bam
```

The extracted BAM file is indexed for efficient access.

Step3:

```
samtools fasta {prefix}.bam > {prefix}.fasta
```

A FASTA file containing the mitochondrial reads is generated.

Step4:

```
mitohifi.py -r {{prefix}.fasta} -f {ref_fa} -g {ref_gb} -t {num_cpus}
-o 1
```

Here, `{bam}` represents the input lrWGS BAM file, `{bai}` the BAM index file, `{locus}` the mitochondrial chromosome (chrM), `{prefix}` a user-defined prefix for output files, `{reads}` the extracted mitochondrial reads FASTA, `{ref_fa}` the reference FASTA file (rCRS), `{ref_gb}` the reference GB file, and `{num_cpus}` the number of CPUs to utilize.

MitoHiFi is executed to generate the mitochondrial assembly using the extracted reads, reference FASTA, reference GB, and specified number of CPUs. Sequences in the `final_mitogenome.fasta` of the output file are utilized for the downstream analysis.

Himito assemblies were generated using the following sequential commands:

Step1:

```
Himito filter -i {lrWGS-bam} -c chrM -m {prefix}_mt.bam -n
{prefix}_numts.bam
```

Himito filters the input lrWGS BAM file, separating mitochondrial reads into `mt.bam` and nuclear mitochondrial sequences (NUMTs) into `numts.bam`.

Step2:

```
Himito build -i {prefix}_mt.bam -k {kmer_size} -r {reference} -o
{sampleid}.{prefix}.gfa
```

Himito constructs a sequence graph using the mitochondrial BAM file with a specified k-mer size and reference, outputting a GFA file.

Step3:

```
Himito asm -g {graph_gfa} -s {sampleid} -o {sampleid}.{prefix}.fasta
```

Himito extracts the primary haplotype sequence from the generated graph, outputting a FASTA file.

Here, `{lrWGS-bam}` represents the same input lrWGS BAM file, `{prefix}` a user-defined prefix, chrM the mitochondrial chromosome identifier, `{kmer_size}` the k-mer size used in graph construction (default 21), `{reference}` the reference FASTA file (rCRS), `{sampleid}` a sample identifier, and `{graph_gfa}` the generated GFA file.

Assembly Comparison and Evaluation

To quantify the accuracy of the MitoHiFi and Himito assemblies, a k-mer-based comparison approach was employed. First, all 100-mers (sequences of length 100 bp) present in the HPRC chrM contigs (truth set) were enumerated. Similarly, all 100-mers present in the MitoHiFi and Himito assemblies were also enumerated. The Jaccard similarity score was then calculated to compare the overlap of k-mers between each assembly and the HPRC truth set. The Jaccard similarity is defined as the size of the intersection divided by the size of the union of the sets of k-mers. Mathematically, Jaccard similarity $J(A, B) = |A \cap B| / |A \cup B|$, where A and B are the sets of k-mers from the assemblies and the truth set, respectively. We use k = 100, 300, 500 to repeat the test. These Jaccard similarity scores were then visualized using boxplots generated with the seaborn library in Python. These boxplots provided a quantitative measure of the similarity between the assemblies generated by each method and the HPRC truth set, allowing for a direct comparison of their performance.

#### In silico Mixture Experiment for Heteroplasmic Variant Calling Evaluation

To evaluate the performance of heteroplasmic variant calling algorithms, we conducted in silico mixture experiments. These experiments were designed to simulate samples with known heteroplasmic variants frequencies, allowing for a quantitative assessment of precision and recall. The workflow can be found here: https://github.com/broadinstitute/Himito/blob/main/wdl/MixSampleExperiment.wdl

Step1: Downsampled to a specific depth

Two HPRC samples, each possessing distinct sets of mtDNA variants, were selected. lrWGS data for these samples were subset to reads aligned to the mitochondrial chromosome (chrM) using

```
samtools view -bhX ~{bam} ~{bai} ~{locus} > ~{prefix}.bam
```

The resulting chrM-aligned read sets were then downsampled to a specific, controlled sequencing depth using GATK toolkit. Deterministic seeds are applied to ensure the reproducibility of this experiment.

```
gatk DownsampleSam -I ~{bam} \
     -O ~{basename}_~{desiredCoverage}x_~{second_fraction}.bam \
     -R 1 \
     -P ~{scalingFactor2} \
     -S ConstantMemory \
     --VALIDATION_STRINGENCY LENIENT \
     --CREATE_INDEX true
```

Step2: Mix the two samples with a predefined ratio

These downsampled BAM files were mixed at predefined ratios, creating mixture samples with known heteroplasmic variant frequencies.

```
samtools merge -f ~{prefix}.merged.bam \
          ~{first_donor_bam} \
          ~{second_donor_bam}
```

Step 3: Creating Truth VCF

The corresponding assembly-based VCF files for the individual donor samples were firstly normed to biallelic files and then merged to generate a truth callset representing the expected variants in the mixed samples.

```
bcftools norm \
        -f ~{reference_fa} \
        -m -both ~{donor_vcf} \
        -O z \
        -o donor.normed.vcf.gz

bcftools merge \
     ~{first_donor_vcf} \
     ~{second_donor_vcf} \
     -O z \
```

```
            -o ~{prefix}.merged.vcf.gz
```

Step 4 Variant calling using Himito, Mitorsaw and GATK-Mutect2

Heteroplasmic variant calling was then performed on the mixed BAM files using three different algorithms: Himito, Mitorsaw, and GATK-Mutect2 in mitochondrial mode.

Himito callset generation:

```
Himito filter -i {merged.bam} -c chrM -m {prefix}_mt.bam -n
{prefix}_numts.bam

Himito build -i {prefix}_mt.bam -k {kmer_size} -r {reference} -o
{sampleid}.{prefix}.gfa

Himito call -g {sampleid}.{prefix}.gfa  -r ~{reference_fa} -k
~{kmer_size} -s ~{sampleid} -o ~{sampleid}.~{prefix}.vcf
```

The `{merged.bam}` is the output mixed BAM file from Step 2. The `{reference_fa}` is the standard rCRS FASTA file. Himito will filter the NUMTs-derived reads first, then output all the mtDNA-derived reads in the `{prefix}_mt.bam`. Then Himito will construct a graph using these mtDNA-derived reads in `{sampleid}.{prefix}.gfa`. The variants are called by using the Himito call module.

The `~{sampleid}.~{prefix}.vcf` is the final Himito output VCF file.

Mitorsaw callset generation:

```
mitorsaw haplotype \
            --reference ~{reference_fasta} \
            --bam ~{merged.bam} \
            --minimum-maf ~{min_af} \
            --output-vcf ~{prefix}.mitorsaw.vcf.gz \
            --output-hap-stats ~{prefix}.mitorsaw.stat
```

Similarly, the `~{merged.bam}` is the output mixed BAM file from Step 2. The `~{reference_fasta}` is the standard rCRS FASTA file. Since there is little documentation about the role of heuristic parameters, we first performed analysis using the default setting for `min_af = 0.1` (Supplementary Figure 4). Mitorsaw showed impressive high precision at all predefined ratios, yet low sensitivities when the heteroplasmic variants frequencies are lower than 0.1. We then performed the experiment using `min_af = 0.01`. The results suggested that Mitorsaw showed higher sensitivity with the cost of precision drops.

Step 5 Callset evaluation using RTG vcfeval

To assess the accuracy of these calls, the generated callsets were compared against the truth callset using RTG vcfeval, a tool for evaluating variant call sets. The VCFs are first normalized to biallelic VCF files.

```
bcftools norm \
        -f ~{reference_fa} \
        -m -both ~{base_vcf} \
        -O z \
        -o ~{base_vcf}.normed.vcf.gz
```

RTG vcfeval provided metrics for precision, recall, and F1 score, enabling a quantitative comparison of the performance of each variant calling method at the simulated heteroplasmic frequencies.

```
rtg format -o rtg_ref ~{reference_fa}
rtg vcfeval \
    -b ~{base_vcf}  \
    -c ~{query_vcf} \
    -o reg \
    -t rtg_ref \
    --squash-ploidy \
    --sample ALT,ALT \
    --vcf-score-field ~{vcf_score_field}
```

#### Benchmarking Long-read Callers using HapMap Mixture Dataset

To evaluate the performance of homoplasmic and heteroplasmic variant calling algorithms in experimental data, we benchmarked using HapMap Mixture PacBio Revio lrWGS downloaded from the SMaHT data portal (https://data.smaht.org/), aligned by pbmm2 (Accession IDs: SMAFI5XI2KWA, SMAFI8IUSZ6W, SMAFIA3UVATP, SMAFIGJIKM6M, SMAFIIUVRY95, SMAFIZH1KWY7)

Step 1:

Subset and DownSampling:

PacBio lrWGS data for these HapMap mixture samples were subset to reads aligned to the mitochondrial chromosome (chrM) using

```
samtools view -bhX ~{bam} ~{bai} ~{locus} > ~{prefix}.bam
```

Using GATK tookit, we downsampled each chrM-aligned read sets to a predefined sequencing depth (10x, 20x, 50x, 80x, 100x, 200x, 300x, 400x, 500x, 600x).

```
gatk DownsampleSam -I ~{bam} \
    -O ~{basename}_~{desiredCoverage}x.bam \
    -R 7 \
    -P ~{scalingFactor} \
    -S ConstantMemory \
    --VALIDATION_STRINGENCY LENIENT \
    --CREATE_INDEX true
```

Step 2:

Variants calling:

We called mitochondrial variants using Himito and Mitosaw respectively.

Himito callset generation:

```
Himito filter -i ~{basename}_~{desiredCoverage}x.bam -c chrM -m
{prefix}_mt.bam -n {prefix}_numts.bam

Himito build -i {prefix}_mt.bam -k 21 -r {reference} -o
{sampleid}.{prefix}.gfa

Himito call -g {sampleid}.{prefix}.gfa -r ~{reference_fa} -k 21 -s
~{sampleid} -o ~{sampleid}.~{prefix}.vcf
```

The `{reference_fa}` is the standard rCRS FASTA file. The `~{sampleid}.~{prefix}.vcf` is the final Himito output VCF file.

Mitorsaw callset generation:

```
mitorsaw haplotype \
            --reference ~{reference_fasta} \
            --bam ~{basename}_~{desiredCoverage}x.bam \
            --minimum-maf 0.01 \
            --output-vcf ~{prefix}.mitorsaw.vcf.gz \
            --output-hap-stats ~{prefix}.mitorsaw.stat
```

The `--minimum-maf` parameter of mitorsaw is set to 0.01 to ensure a balanced comparison with Himito.

Step 3:

Evaluation:

We then used RTG vcfeval to evaluate the performance against the truth set. The truth set is curated by the SMaHT consortium and is available in https://data.smaht.org/.

```
rtg format -o rtg_ref ~{reference_fa}
rtg vcfeval \
    -b ~{base_vcf}  \
    -c ~{query_vcf} \
    -o reg \
    -t rtg_ref \
    --squash-ploidy \
    --sample ALT,ALT \
    --vcf-score-field ~{vcf_score_field}
```

Step 4:

Visualization

We plot the precision, recall and F1 scores at different coverage using python seaborn.

```
sns.boxplot(df_summary, x = "Depth", y = "Score", hue = "Type")
```

We further optimized the parameter of Himito for ONT long read assembly and variant calling. We performed the same analysis using 5 ONT samples in the HapMap mixture dataset: SMAFIBT8WCXG, SMAFIF7CG4YZ, SMAFIPF11BMW, SMAFIQHVZFJG, SMAFIWI78FL3. Their chrM mean depth is 447.166, 382.183, 369.203, 5511.97, 7881.3 respectively. We downsampled these BAMs into 10x, 20x, 50x, 80x, 100x, 200x, 300x. The evaluation results are in Supplementary Figure 17.

#### Benchmarking against short read sequence data

##### Evaluation between short-read and long-read data using HapMap Mixture

To compare mitochondrial variant calling performance between lrWGS and short-read whole genome sequencing (srWGS) data modalities, we downloaded HapMap Mixture srWGS data from the SMaHT data portal (https://data.smaht.org/), which had been aligned to the GRCh38 using BWA-MEM. Unique accession IDs distinguish the srWGS HapMap Mixture BAM files from sequencing lanes processed by four Genome Characterization Centers (BCM, WashU, NYGC, Broad): SMAFI3EQANWU, SMAFIB1SYCDQ, SMAFIVF1NGTJ, SMAFIXLDO5QN, SMAFI9RHZRDY, SMAFIWYI2RF6, SMAFI1LJ41ED, SMAFIG7OXW7O, SMAFIICOPTLS, SMAFIPERCVSF, SMAFIR37VX6A. HapMap Mixture srWGS data from each GCC were subset to reads aligned to chrM using samtools as previously described.

The resulting srWGS chrM-aligned read sets were then downsampled to 600X using the same method as the lrWGS data.

```
gatk DownsampleSam \
        -I ~{bam} \
        -O ~{OUT_DIR}/~{basename}_~{desiredCoverage}x.bam" \
        -R 7 \
        -P ~{scalingFactor} \
        -S ConstantMemory \
        --VALIDATION_STRINGENCY LENIENT \
        --CREATE_INDEX true
```

Variant calls from HapMap srWGS data were generated using three short-read mtDNA callers: Mutect2 in mitochondrial mode, mtDNA-Server (mutserve), and MToolBox. The following commands were used for each method:

```
gatk Mutect2 -R ~{reference_fasta} -L chrM --mitochondria-mode -I
~{srWGS-bam} -O ~{prefix}.mutect2.vcf.gz

nextflow run genepi/mtdna-server-2 -r v2.1.16 -c ~{config_file} -
profile docker

MToolBox.sh -i ~{config_file}
```

Subsequently, both the long-read variant calls from Himito and Mitorsaw and the short-read calls from Mutect2 mito-mode, mtDNA-Server, and MToolBox were evaluated using RTG vcfeval.

```
rtg vcfeval -T 10 \
      --template ~{sdf_hg38} \
      --baseline ~{truth_hg38} \
      --calls ~{vcf} \
      --sample ~{HapMap_Mixture,~{calls_sample}} \
      --squash-ploidy \
      --evaluation-regions ~{bed} \
      --all-records \
      --output ~{SNV_~{caller}_~{sample_id}_total}
```

##### Concordance between short-read and long-read callset

To further validate the long-read based mitochondrial variant calling, we compared our results with those derived from paired short-read whole genome sequencing (srWGS) data from the HPRC samples. High-confidence variant calls from srWGS data were established using GATK Mutect2 in mitochondrial mode. Specifically, the following command was used:

```
      gatk Mutect2 -R ~{reference_fasta} -L chrM --mitochondria-mode -I ~{srWGS-bam}
-O ~{prefix}.mutect2.vcf.gz
```

Here, `{srWGS-bam}` represents the full-coverage srWGS data, and `{reference_fasta}` is the rCRS FASTA file. The resulting high-confidence variant calls were stored in `~{prefix}.mutect2.vcf.gz` and served as our truth set for comparison.

Long-read sequencing data were downsampled to specific sequencing depths. Variant calling on these downsampled long-read datasets was performed using the Himito pipeline.

```
Himito filter -i {bam} -c chrM -m {prefix}_mt.bam -n {prefix}_numts.bam

Himito build -i {prefix}_mt.bam -k {kmer_size} -r {reference} -o
{sampleid}.{prefix}.gfa

Himito call -g {sampleid}.{prefix}.gfa -r ~{reference_fa} -k ~{kmer_size} -s
~{sampleid} -o ~{sampleid}.~{prefix}.vcf
```

The `{bam}` is the subset Bam from lrWGS data. The `{reference_fa}` is the standard rCRS FASTA file. Himito will filter the NUMTs-derived reads first, then output all the mtDNA-derived reads in the `{prefix}_mt.bam`. Then Himito will construct a graph using these mtDNA-derived reads in `{sampleid}.{prefix}.gfa`. The variants are called by using the Himito call module. The `~{sampleid}.~{prefix}.vcf` is the final Himito output VCF file.

Subsequently, both the Himito-derived variant calls and the GATK Mutect2 calls were evaluated using RTG vcfeval. This allowed for a quantitative comparison of the callsets and an assessment of the performance of Himito against the established short-read benchmark.

```
rtg vcfeval \
           -b ~{base_vcf}  \
           -c ~{query_vcf} \
           -o reg \
           -t rtg_ref \
           --squash-ploidy \
           --sample ALT,ALT \
           --vcf-score-field ~{vcf_score_field}
```

For all true positive (TP) calls, the R-squared value was calculated. This value represents the correlation between the allele frequency pairs derived from the Himito long-read callset and the GATK Mutect2 short-read callset, providing a measure of concordance between the two techniques. Higher R-squared values indicate a stronger agreement in allele frequencies, suggesting that Himito accurately reflects the variant frequencies identified by the high-confidence short-read method.

#### Methylation Calling Evaluation Methods

Himito's methyl calling functionality was benchmarked against PbCpG tools in count mode. Himito's methylation calls were generated using the command:

```
Himito methyl -g ~{graph_gfa} \
              -p ~{min_prob} \
              -b ~{bam} \
              -o ~{sampleid}.~{prefix}.bed
```

where `~{graph_gfa}` represents the Himito-constructed sequence graph, `~{min_prob}` is the minimum probability threshold for methylation calling. We used the default setting 0.5. `~{bam}` is the input BAM file, and `~{sampleid}.~{prefix}.bed` is the output BED file containing Himito's methylation calls.

PbCpG methylation calls were generated using the command:

```
/pb-CpG-tools-v2.3.2-x86_64-unknown-linux-gnu/bin/aligned_bam_to_cpg_scores \
                --bam ~{bam} \
                --output-prefix ~{outputprefix} \
                --pileup-mode count \
                --threads ~{num_threads}
```

where `~{bam}` is the input BAM file, `~{outputprefix}` is the prefix for output files, and `~{num_threads}` is the number of threads to use.

A key difference was noted in the coordinate systems used: Himito Methyl BED files utilize 1-based coordinates, while PbCpG calls are reported in 0-based coordinates. To ensure accurate comparison, coordinate adjustments were performed to align the data from both methods. Subsequently, the R-squared value was calculated for the modification scores of each corresponding CpG site pair between the Himito and PbCpG results. This R-squared value represents the correlation between the modification scores obtained from the two methods, serving as a metric to evaluate the concordance of methylation calling between Himito and PbCpG tools.

### AoU v8 Callset Analysis

#### Sample Depth Analysis

Himito was used to analyze genetic variations in All of Us v8 long read samples. This release includes 2517 PacBio and 324 ONT samples, with one individual sequenced on both PacBio Revio and Sequel platforms.

Initially, the sequencing depth of each sample on the mitochondrial chromosome (chrM) was evaluated. Samples with an average depth below 10x were removed. As a result, 2422 PacBio samples met the depth requirement for further analysis.

The mean coverage on chrM for AoU v8 PacBio samples was calculated using the following samtools command:

```
samtools depth \
    -a \
    -r ~{region} \
    ~{bam} > ~{sampleid}.depth.txt
```

Here, `~{bam}` represents the lrWGS BAM file and `~{region}` specifies "chrM." This command outputs depth at all positions on chrM. Samples with mean depths below 10x were excluded.

These text files were then loaded and aggregated using the Python pandas package. The mean coverage and standard deviation for each site were calculated. Finally, Seaborn was used to generate the corresponding coverage plot.

#### Variant Calling and Data Processing for AoU v8 PacBio Samples

For each PacBio long-read sequencing sample from the All of Us v8 release, single-sample VCF files were generated using the Himito pipeline. The Himito workflow version available at https://github.com/broadinstitute/Himito/blob/main/wdl/Himito_call.wdl was employed with default parameters: k-mer size (k) set to 21, minimum allele count (minimal-ac) set to 1, and variant allele frequency (vaf-threshold) set to 0.01.

The variant calling process involved the following sequential steps:

1. **Read Filtering:** Reads from the lrWGS BAM file were filtered using the `Himito filter` module. Reads aligned to the mitochondrial chromosome (chrM) were extracted and designated as `prefix_mt.bam`, while nuclear mitochondrial sequences (NUMTs) were separated into `prefix_numts.bam`. The methylation filtering threshold is 0.2. The command used was:

```
Himito filter -i {bam} -c chrM -m {prefix}_mt.bam -n {prefix}_numts.bam
```

2. **Graph Construction:** A sequence graph was constructed using the `Himito build` module. The mitochondrial reads `prefix_mt.bam` generated by `Himito filter` step were utilized with a specified k-mer size (`kmer_size`) and reference sequence (`reference`). The resulting graph was outputted as a GFA file (`sampleid.prefix.gfa`). The command used was:

```
Himito build -i {prefix}_mt.bam -k {kmer_size} -r {reference} -o
{sampleid}.{prefix}.gfa
```

3. **Variant Calling:** Variants were called from the generated graph using the `Himito call` module. The graph file generated by `Himito build`, the rCRS reference FASTA file (`reference_fa`), k-mer size (`kmer_size`), sample identifier (`sampleid`), and output VCF file (`sampleid.prefix.vcf`) were specified. The command used was:

```
Himito call -g {sampleid}.{prefix}.gfa -r ~{reference_fa} -k ~{kmer_size} -s
~{sampleid} -o ~{sampleid}.~{prefix}.vcf
```

**Merging and Genotype Matrix Generation**

Following single-sample variant calling, mitochondrial variants from all samples were merged using `bcftools merge`. The following commands were employed:

```
bcftools merge \
        --threads ~{threads_num} \
```

```
                --force-single \
                --merge none \
                -l merge.txt \
                -O z \
                -o ~{pref}.AllSamples.vcf.gz
bcftools index --threads ~{threads_num} -t ~{pref}.AllSamples.vcf.gz
```

A genotype matrix was then extracted from the merged callset. In this matrix, a value of 1 represented the presence of a variant in a specific individual (regardless of allele frequency), and 0 indicated the absence of the variant.

**Principal Component Analysis (PCA)**

Principal Component Analysis (PCA) was performed based on the extracted genotype matrix. The PCA was conducted using the `sklearn` package in Python. Samples are color-coded based on haplogroup information determined by haplogrep v2.4.0.

#### Haplogroup Classification

Haplogroup determination for each individual was performed using Haplogrep2 (v2.4.0). The specific command line employed is as follows:

```
haplogrep classify --in ~{vcf} --format vcf --output ~{prefix}.haplogroups.txt
```

The merged multi-sample VCF, produced by `bcftools merge`, is denoted as `~{vcf}`. Haplogroup classification results are outputted as a text file, providing each individual's classified haplogroup. The first two digits of the haplogroup are used for this classification. PCA analysis results are color-coded based on haplogroup information.

#### Functional Annotation using VEP

Functional annotation of mitochondrial variants detected by Himito was performed using the Variant Effect Predictor (VEP). The command line employed here is:

```
vep    --input_file ~{vcf} \
       --dir_cache ~{cache_directory}    \
       --fasta ~{reference_fasta} \
       --force_overwrite   \
       --hgvs       \
       --hgvsg      \
       --offline    \
       --no_escape  \
       --protein    \
       --symbol     \
       --tab        \
       --output_file ~{prefix}.vep.annotation.txt
```

where `--input_file ~{vcf}` specifies the input VCF file; `--dir_cache ~{cache_directory}` points to the directory for cached data, the reference FASTA file location is specified by `--fasta ~{reference_fasta}`, and options to force overwrite existing files `--force_overwrite`, enable HGVS output `--hgvs --hgvsg`, run in offline mode `--offline`, include protein information `--protein`, include gene symbols `--symbol`, output in tab-delimited format `--tab`, and specify the output file `--output_file ~{prefix}.vep.annotation.txt`.

VEP predicted the functional consequences of each variant, classifying them as synonymous, missense, frameshift (leading to amino acid sequence alterations), or stop gained (introducing a premature stop codon). These variants were further categorized by their predicted consequences, and the heteroplasmic frequencies of each catalog were visualized using the Seaborn library. Additionally, VEP evaluated the potential impact of each variant, classifying them as High, Moderate, or Low.

#### Pathogenic Variants analysis

MitoMap, a database of human mitochondrial DNA variation, catalogs reported pathogenic mutations and polymorphisms. It is a valuable resource for researchers studying mitochondrial genetics and disease. Pathogenic variants in mtDNA were obtained from https://www.mitomap.org/foswiki/bin/view/MITOMAP/ConfirmedMutations. The Himito callset was then cross-referenced with pathogenic variants from the MitoMap database. The identified pathogenic variants were extracted, and their presence across individuals was subsequently analyzed using a seaborn heatmap.

##### Length Heteroplasmic Variants Analysis

`Himito call` will output an occurrence matrix of variants versus reads presented in the graph. We subset the variants to three length heteroplasmic loci: chrM:302-316 (ACCCCCCCTCCCCCG), chrM:567-574 (ACCCCCCA), and chrM:16174-16195 (CAAAACCCCCTCCCCA). For each read, we reconstruct the haplotype by replacing the reference alleles with the alternative alleles reported in each loci. We then calculate the number of length heteroplasmy haplotypes per sample and the joint haplotypes across each pair of locus. We further aggregate the reads from more than 2000 PacBio samples and construct an occurrence matrix for each length heteroplasmy allele in 229985 reads. For each pair of alleles, we performed Fisher's Exact test to evaluate the association between the occurrence of two alleles. The Fisher's Exact test is performed by using `fisher_exact` module in `scipy.stats`. The pair of alleles with p values < 0.001 were retained and are plotted by using `seaborn.heatmap()`.

#### AoU v8 Pangenome Construction and Analysis

##### AoU v8 Pangenome Construction

**Primary Assembly Generation:** Individual primary mitochondrial genome assemblies were constructed for each sample within the All of Us v8 PacBio long-read sequencing cohort using the Himito pipeline. Briefly, raw long-read whole-genome sequencing (lrWGS) BAM files were processed to isolate mitochondrial DNA (mtDNA)-derived reads and exclude reads originating from nuclear mitochondrial sequences (NUMTs). This was achieved using the `Himito filter` module with the command:

```
Himito filter -i {lrWGS-bam} -c chrM -m {prefix}_mt.bam -n
{prefix}_numts.bam
```

where `lrWGS-bam` denotes the input lrWGS BAM file, `chrM` specifies the mitochondrial chromosome identifier, and `{prefix}` is a user-defined prefix for output files. The resulting `{prefix}_mt.bam` file contained mtDNA-derived reads, while `{prefix}_numts.bam` stored NUMTs-derived reads.

Subsequently, a sequence graph was constructed for each individual sample using the `Himito build` module:

```
Himito build -i {prefix}_mt.bam -k {kmer_size} -r {reference} -o
{sampleid}.{prefix}.gfa
```

Here, `{prefix}_mt.bam` is the mtDNA-derived read file generated by `Himito filter`, `-k {kmer_size}` sets the k-mer size to 21 bp, `{reference}` specifies the rCRS reference FASTA file, and `{sampleid}` is a sample identifier. The output `{sampleid}.{prefix}.gfa` file represents the individual's mitochondrial genome sequence graph. Primary haplotype sequences were extracted from each graph using the `Himito asm` module:

```
Himito asm -g {graph_gfa} -s {sampleid} -o {sampleid}.{prefix}.fasta
```

where `{graph_gfa}` is the individual's sequence graph file and `{sampleid}.{prefix}.fasta` stores the resulting primary mitochondrial haplotype assembly.

**Pangenome Graph Construction:** Individual primary assembly FASTA files were concatenated into a single multi-sequence file (`all.fasta`) using the `cat` command:

```
cat fasta1 fasta2 … > all.fasta
```

This `all.fasta` file, representing the complete set of individual mitochondrial assemblies, was then used as input to the `Himito build` module to construct a population-scale pangenome graph:

```
Himito build -i all.fasta \
             -k ~{kmer_size} \
             -r ~{reference} \
```

```
            -o population.gfa
```

The `-k ~{kmer_size}` set parameter of the k-mer size at 21 bp, `{reference}` again specifies the rCRS reference FASTA, and `population.gfa` is the output file containing the population-scale pangenome graph representing mitochondrial genome diversity across the AoU v8 cohort.

Because Himito-generated sequences all begin at the same mitochondrial genome location, specifically the initial anchor (the first unique k-mer in the rCRS genome), the population sequence graph initially forms as linear, not circular. To create a circular pangenome graph, we manually join the sequences preceding the first anchor and following the last anchor for each haplotype. This step allows for a precise evaluation of sequence variability within the D-loop region.

**Hierarchical clustering of the presence and absence of graph vertices in each sample**

A presence/absence matrix of graph vertices in each sample was created to study genomic variation across the population. Using the Seaborn Python library, this matrix was then subjected to hierarchical clustering. The command is:

```
sns.clustermap(
    df,
    method='complete',
    col_colors=sample_groups.map(group_colors),
    figsize=(12, 10),
    cmap = "Blues"
)
```

Where `df` is the binary matrix representing the presence or absence of the graph vertices in each sample; each sample is color-coded by their haplogroups.

#### Haplotype Number determination

We developed a novel approach to estimate the haplotype diversities within certain genomic regions in the mitochondrial genome. This method began by establishing a coordinate framework centered on anchor sequences. Each anchor was defined as a unique k-mer within the standard reference mitochondrial genome, intrinsically linked to a specific reference genomic coordinate. The anchors functioned as a sequence-based coordinate system, allowing for the accurate definition of genomic regions. This effectively linked graph vertices to specific genomic intervals by the bounding anchor sequences. For each genomic region defined by the reference coordinates, we first identify all the graph vertices overlapping with this genomic interval using anchor-based coordinates. Utilizing the inherent graph connectivities, we reconstructed subgraphs that represented the sequence diversities within the specific genomic region. Enumeration of distinct haplotype sequences within these subgraphs was achieved through graph traversal, ensuring that every possible combination of variations supported by a minimal number of long reads was identified and recorded. To elucidate the sequence differences between haplotypes, multiple sequence alignments (MSA) were performed. The `ClustalW v2.1` was employed for MSA analysis. The command line employed is shown here:

```
clustalw2 -infile=input_clustal.fa
```

The `input_clustal.fa` includes all the haplotypes extracted from a subgraph.

Visualization of these alignments was conducted using the python package MSAplot, available at https://github.com/mourisl/MSAplot.

To evaluate the haplotype diversity across the whole mitochondrial genome, we divided the entire mitochondrial genome into non-overlapping 1kb windows. Within each of these windows, we calculated the number of distinct haplotypes present, providing a quantitative measure of local diversity. These counts were then visualized using a circos plot, which was generated using the Python package `pyCirclize 1.9.1` (https://github.com/moshi4/pyCirclize), allowing for an effective and holistic view of the haplotype number distribution.

#### Sequence Diversity Score Calculation

The gene intervals used for this analysis were obtained from the GFF3 file of the standard rCRS reference genome, which was downloaded directly from the National Center for

Biotechnology Information (NCBI). This file provides feature annotation and genomic coordinates for mitochondrial genome. For each defined gene interval, a subgraph was extracted from the AoU v8 pangenome using the same method described above. Within these extracted gene subgraphs, the distinct haplotype sequences were extracted by graph traversal. To quantify genetic diversity, we calculated the count of these distinct haplotype sequences for each gene. This raw count was then normalized by dividing by the length of the corresponding gene. This normalized score serves as a quantitative measure representing the level of diversity observed within each individual gene of the mitochondrial genome. A higher score indicates greater diversity of this gene within the population.

#### Detailed Explanation of Methylation Data Processing

Conventionally, the core methylation information captured for each read includes the relative position of each CpG site (recorded in the MM field) and the corresponding methylation likelihood (recorded in the ML field). This likelihood is represented as an integer value ranging from 0 to 255, where a higher value indicates a greater probability of methylation. To obtain a normalized methylation likelihood for each CpG site, the raw ML value is divided by the maximum possible value, 255.

Subsequently, these individual CpG methylation likelihoods are mapped onto the Himito single-sample graph. The normalized methylation likelihood is then annotated to the corresponding vertex within a "methyl" field. This field stores methylation information for each vertex in a structured format. Specifically, for each read (e.g., "read1", "read2"), the "methyl" field contains a list of paired values: the offset or position of the CpG within the vertex and its associated normalized likelihood. For instance, the format is represented as:

```
{"methyl":{"read1":[[offset,likelihood],[offset,likelihood],...]
,"read2":[[offset,likelihood],[offset,likelihood],...]}}.
```

To gain insights at the haplotype level, the single-read CpG methylation data is further aggregated. Reads that have been assigned to the same haplotype are aligned, and for each CpG site within that haplotype, the modification rate is calculated. This aggregation involves considering the methylation status of the same CpG site across multiple reads belonging to the same haplotype.

A heuristic threshold (t), with a default value of 0.5, is applied to determine whether a CpG site is considered methylated or unmethylated for the purpose of calculating the modification rate. If the normalized methylation likelihood for a CpG site is greater than the threshold (t), it is counted as modified. Conversely, if the likelihood is less than 1-t, the CpG is considered unmodified. CpG sites with likelihood values falling within the intermediate range (between 1-t and t) are excluded from the analysis to ensure a more confident determination of methylation status. The modification rate for each CpG site on a haplotype is then calculated by dividing the number of modified CpG observations by the total number of CpG observations included in the analysis (i.e., those with likelihoods outside the excluded range).

Finally, the aggregated CpG methylation information is mapped back to both the reference genome coordinates and the coordinates of the assembled genome. This allows for the integration of methylation data with existing genomic annotations and the characterization of methylation patterns within the specific context of the assembled genome.

The processed methylation information is reported in standard BED files, providing a comprehensive record of CpG methylation across the genome. Additionally, a CSV file is generated, specifically focusing on the read-level methylation likelihood along the major haplotype, offering a more detailed view of methylation patterns at the individual read level. Analysis can be done based on this methylation matrix to study interesting biological features such as the strand-bias methylation patterns in the mitochondrial genome.

#### Methylation Analysis in the All of Us (AoU) v8 Long-read Release Using PacBio and ONT

For each individual sample, BED files containing methylation information were generated using the `Himito methyl`. Subsequently, modification rate matrices, specific to each sequencing technology (PacBio and ONT), were constructed to quantify the extent of methylation across the genome.

To gain an initial overview of the dataset, a Principal Component Analysis (PCA) was performed on the 41 paired PacBio and ONT samples (Supplementary Figure 7). The PCA plot allowed for the exploration of the primary sources of variation within the data, highlighting differences arising from the sequencing platforms.

Further analysis focused on the distribution of modification rates observed in both PacBio and ONT datasets (Supplementary Figure 8). Based on these distributions, customized methylation rate thresholds were established to categorize individual CpG sites as either hyper-methylated or hypo-methylated. A threshold of 0.3 was chosen for PacBio samples, while a lower threshold of 0.1 was applied to ONT samples. Applying these thresholds, the distribution of hyper-methylated CpG sites across the entire cohort of PacBio and ONT samples was visualized to provide a comprehensive view of methylation patterns in the mitochondrial genome (Supplementary Figure 9).

#### Age-related Analysis

To explore the relationship between age and SNV number and methylation, the number of SNVs and hyper-methylated CpG sites was quantified within distinct age groups using the previously defined technology-specific thresholds. The samples in the AoU v8 cohort were grouped by their reported age. The number of low-frequent SNVs and hyper-methylated CpG sites were summarized in each bin with a sliding window of 10 years. We calculated the number of SNVs (1-10%) and methylated sites in each individual sample by using Himito `call` and `methyl`. The SNVs are further strategies by their types (ref to alt) and each type of variants and their number in each age group were plotted. The CpG methylation likelihood threshold is set as 0.5, and the modification rate of each CpG site was reported by Himito `methyl`, and further aggregated with age groups. We further use a linear regression model (Ordinary least squares) to model the trend of SNVs and methylation increasing with age. We set up a linear model (SNV~ methylation + age ) using a python package *sklearn*. We calculated the p-value of each variable for the signal and found that methylation has a significant negative correlation with low frequent SNVs. This analysis aims to identify potential age-associated changes in DNA low-frequent SNVs and methylation profiles within the AoU v8 cohort, providing insights into the role of epigenetics in aging.

### Supplementary Figures

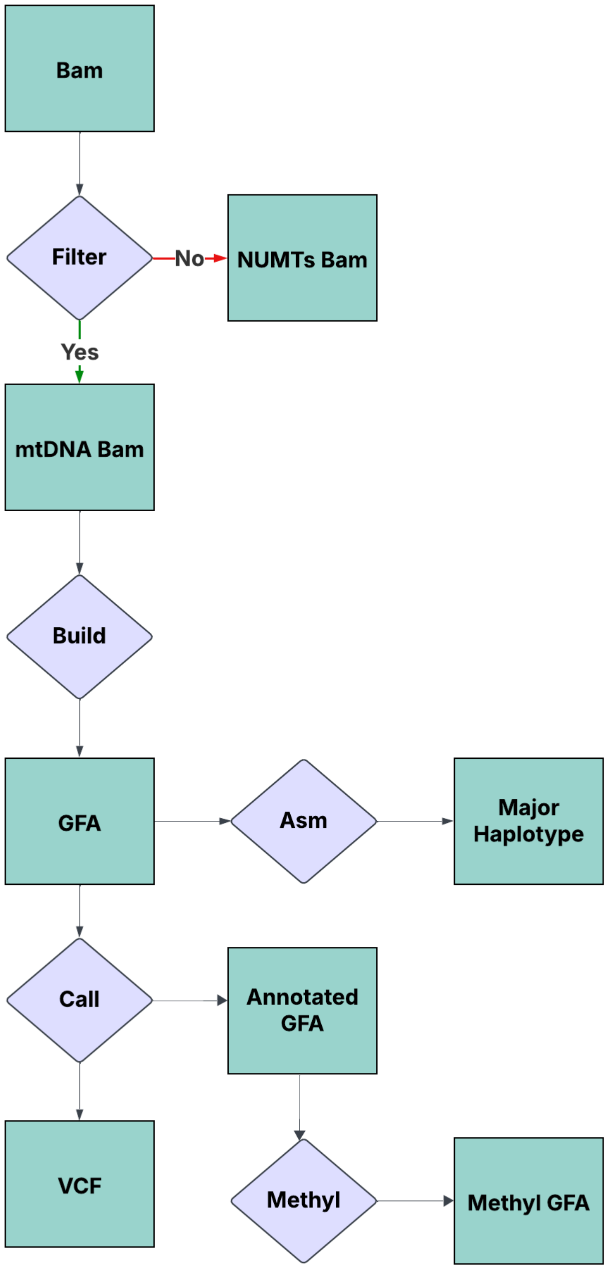

Supplementary Figure 1 Workflow for Himito.

Himito comprises 5 key modules for comprehensive mitochondrial genome analysis. Filter takes lrWGS as input to classify reads as mtDNA-derived and NUMTs-derived reads. It aims to remove NUMTs-derived reads from the downstream analysis. Build constructs a sequence graph to represent the diversity of mtDNAs. Asm takes the sequence graph as the input and extracts primary haplotypes from the graph. Call performs pair-wise alignment between parallel paths and aims to call both homoplasmic and heteroplasmic variants from the graph. Methyl annotates the methylation signals to the graph and aggregates the methylation signals along the major haplotype.

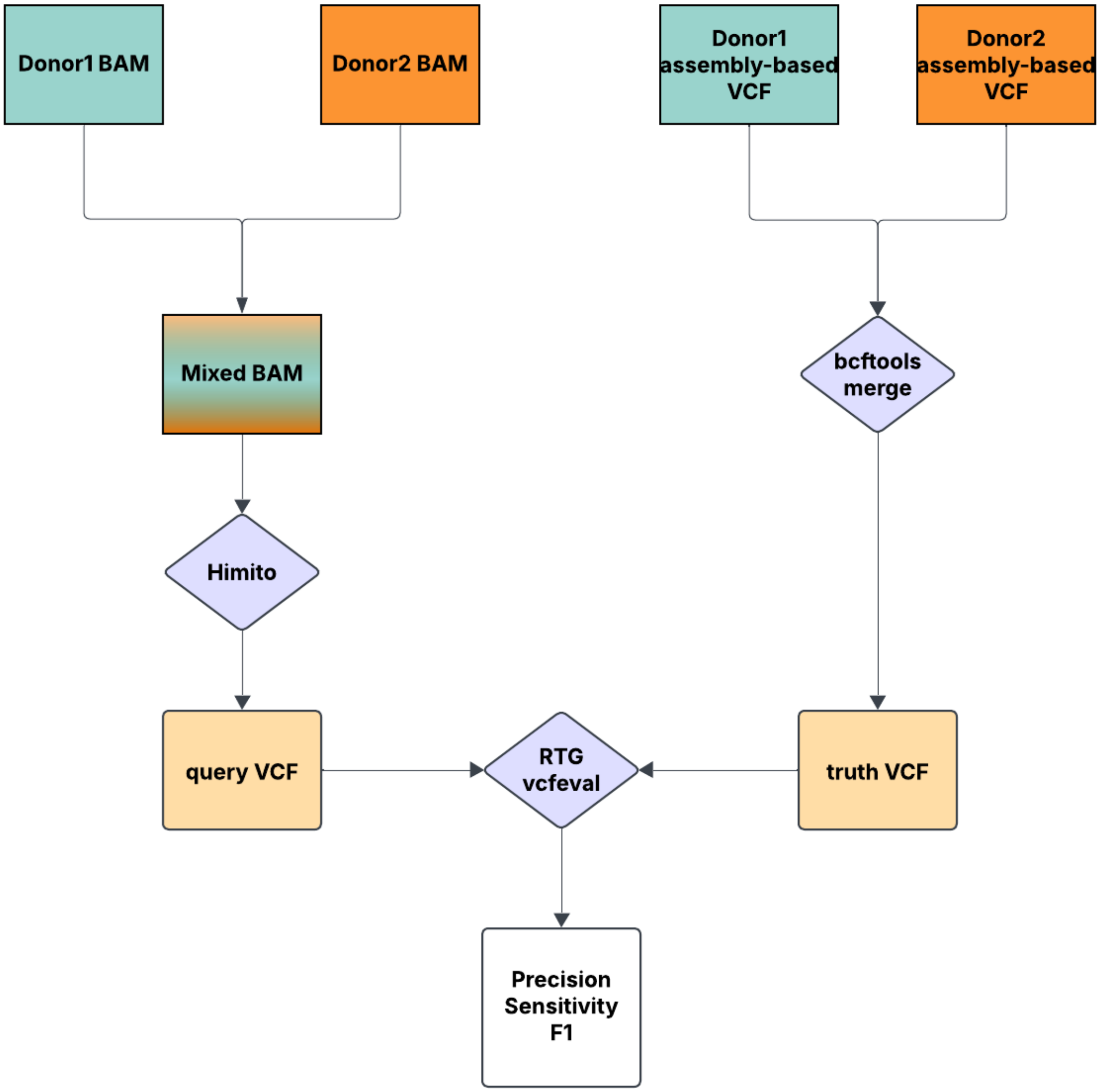


Supplementary Figure 2 In silico mixture experiment for heteroplasmic calling evaluation.

In silico mixture experiments were designed and performed to evaluate the heteroplasmic variant calling performance at predefined frequencies. The performance of the variant caller on the mixed sample was assessed by comparing the called variants against the truth set using the RTG vcfeval tool, allowing for the quantification of precision and recall at the simulated heteroplasmic frequency.

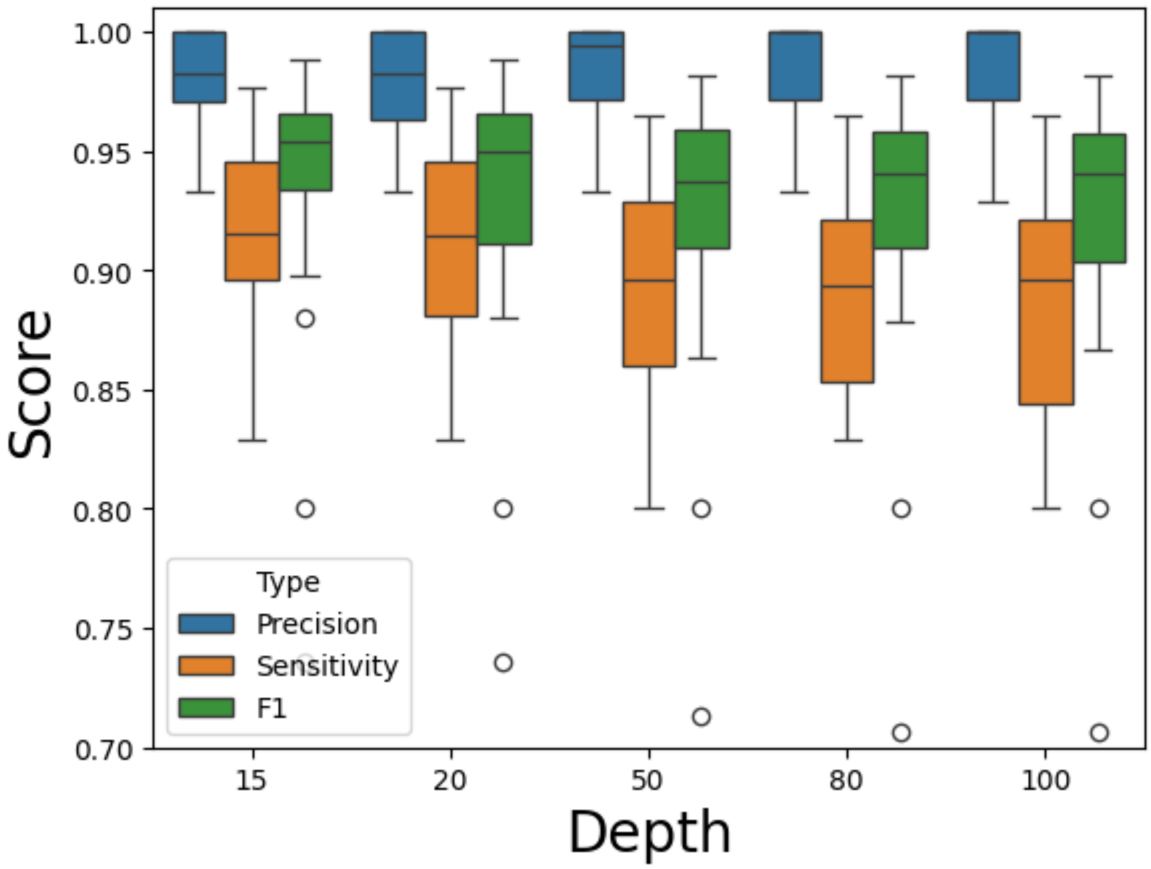


Supplementary Figure 3 Homoplasmic calling evaluation.

The homoplasmic variants (HF>95%) in the Himito callset are compared with the HPRC assembly-based truth callset (generated by minigraph-cactus). The HPRC truth callset comprises both homoplasmic variants and major heteroplasmic variants. Using a heteroplasmy frequency (HF) cutoff of 95% might miss some major heteroplasmic variants in the assembly-based truth callset, potentially leading to reduced sensitivity.

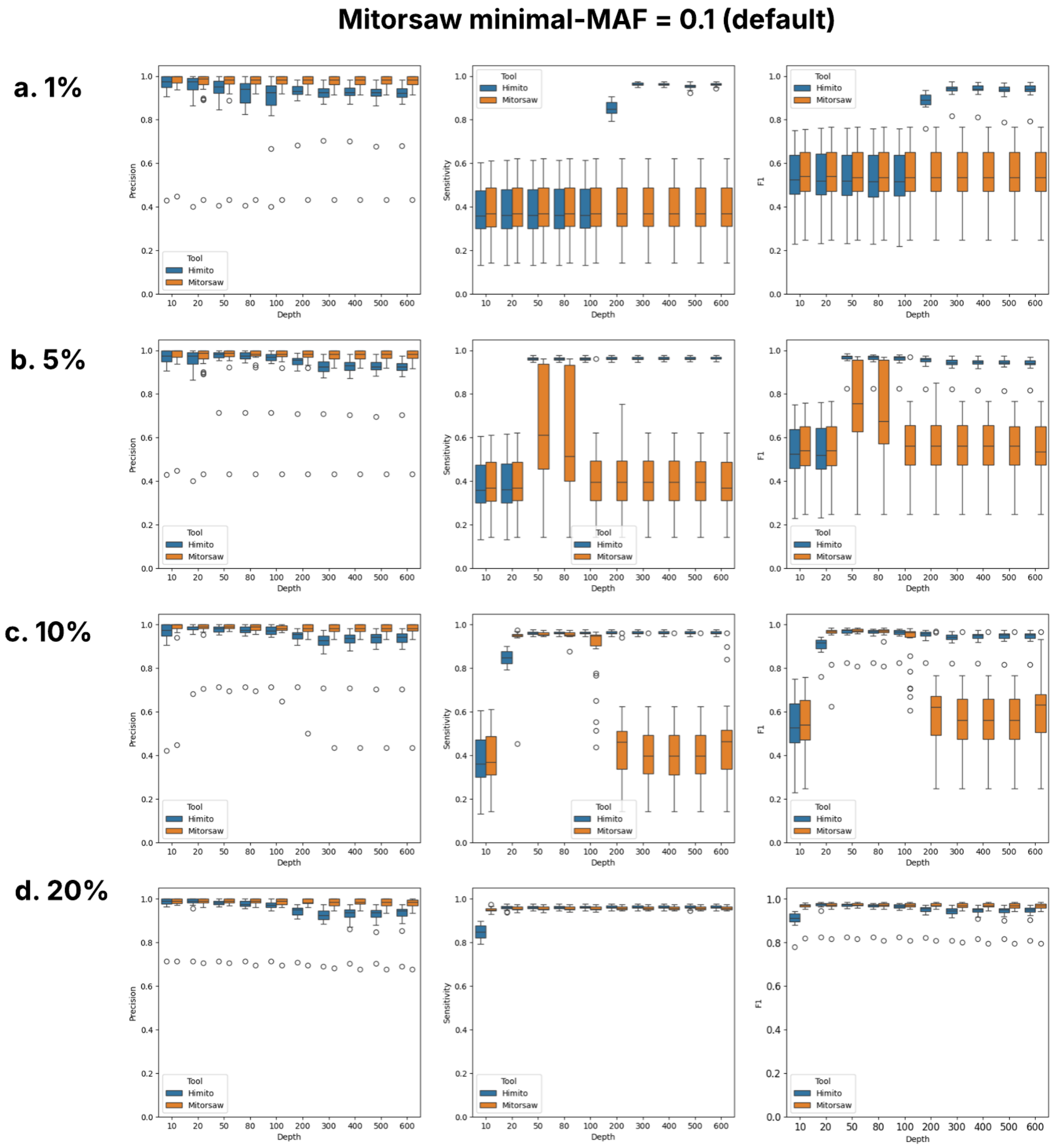


Supplementary Figure 4 Variant calling benchmark against Mitorsaw at default parameters (Mitorsaw `minimal-MAF = 0.1`, Himito `vaf-threshold = 0.01`).

In silico experiments evaluated heteroplasmic frequencies at 0.01 (a), 0.05 (c), 0.1 (d), and 0.2 (e). The precision, sensitivity, and F1 score were calculated for each frequency. Downsampling seed -R = 1.

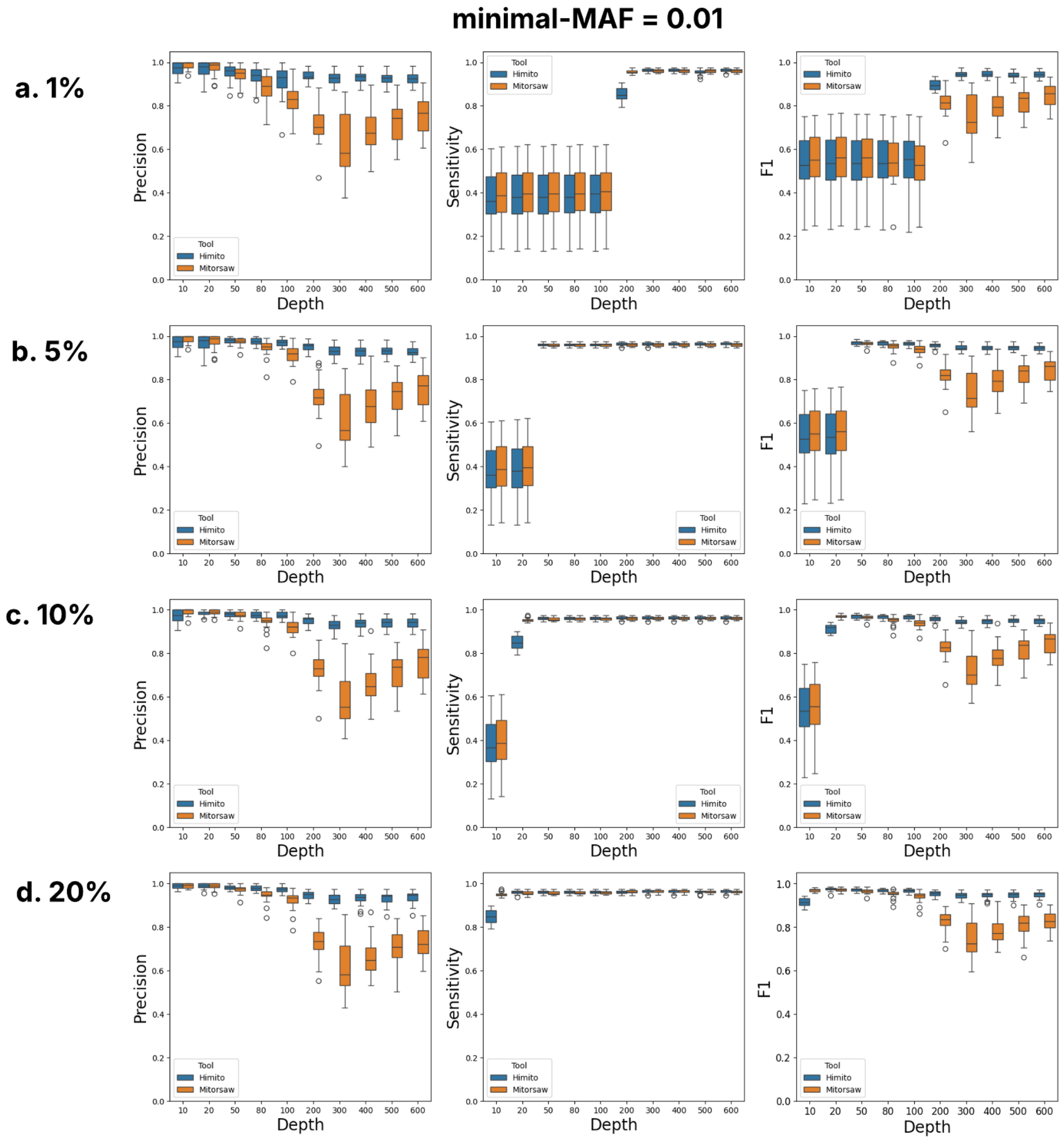


Supplementary Figure 5 Mitorsaw and Himito comparison with the same detection limits (Mitorsaw `minimum-maf = 0.01`, Himito `vaf-threshold = 0.01`).

By changing minimal-MAF from 0.1 to 0.01, Mitorsaw's increased sensitivity comes at the expense of reduced precision. Using in silico experiments, heteroplasmic frequencies at 0.01 (a), 0.05 (c), 0.1 (d), and 0.2 (e) were evaluated. The precision, sensitivity, and F1 score were calculated for each frequency. Downsampling seed -R = 1

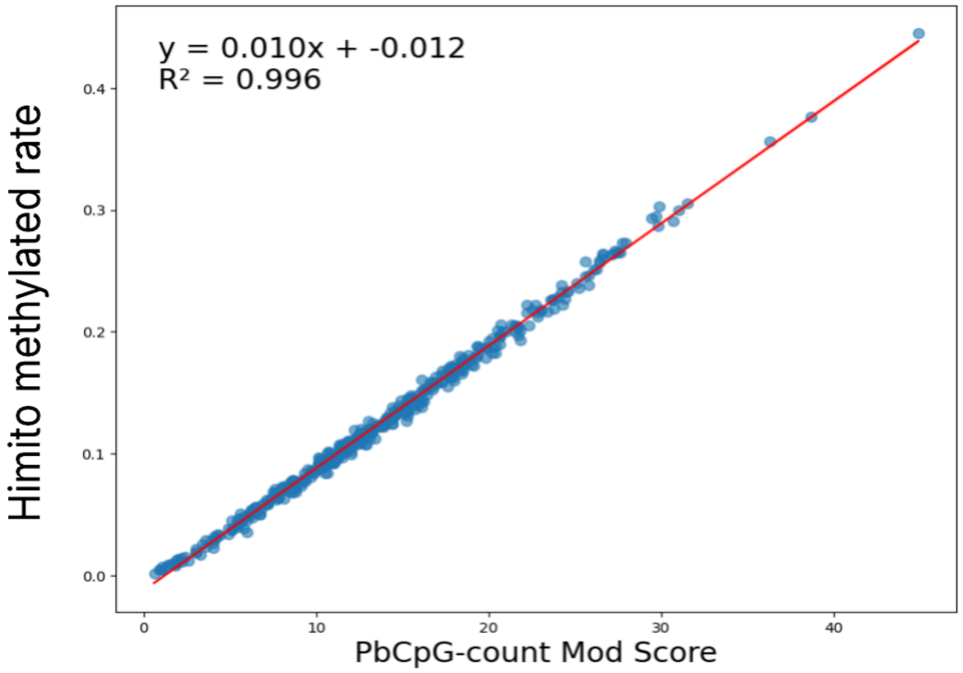


Supplementary Figure 6, Benchmark of Himito Methyl module against PbCpG-count Mode.

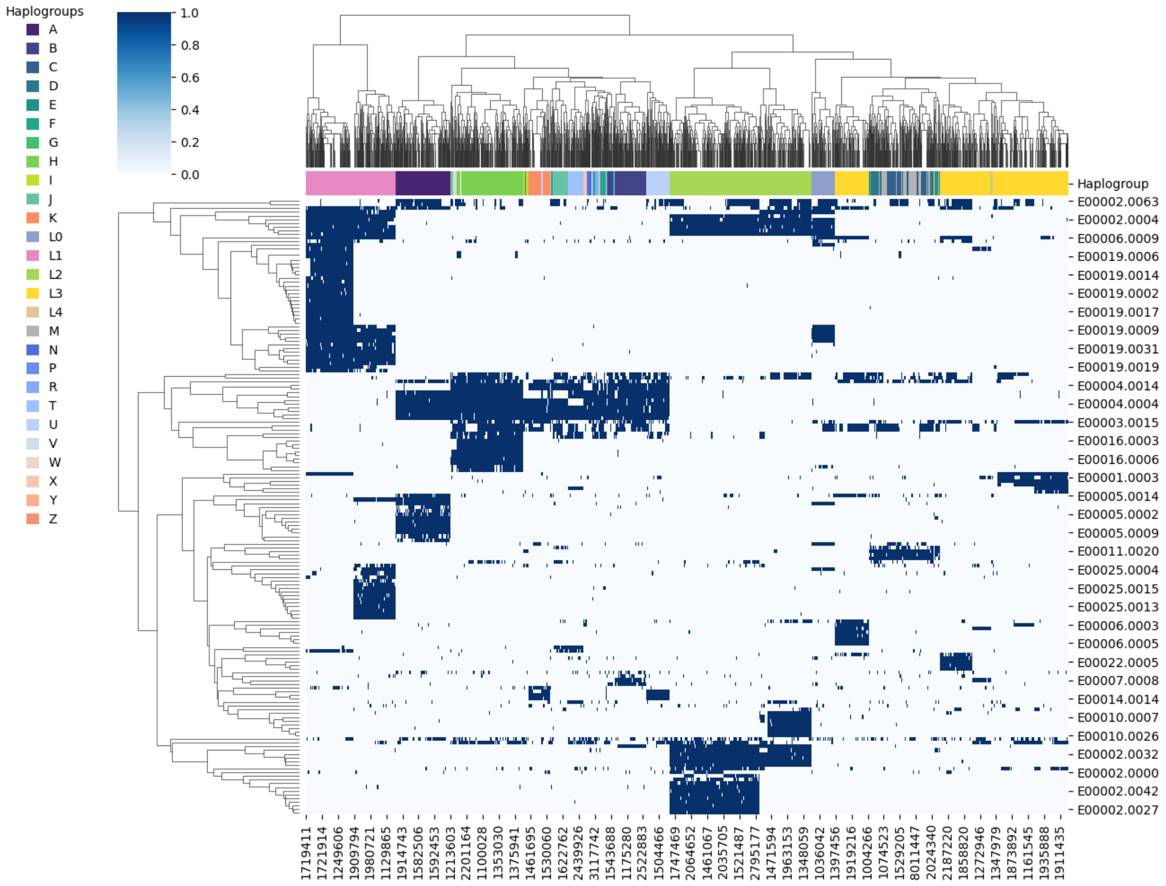


Supplementary Figure 7 Hierarchical clustering of the graph vertices presence-absence matrix in each assembly.

A binary matrix was created to represent the presence or absence of each graph vertex within individual assemblies. Hierarchical clustering of this matrix revealed an automatic grouping of samples according to their haplogroups. Graph vertices with coverage below 100 or above 1000 were excluded from this analysis (sequences less than 100 individuals supported or shared by more than 1000 individuals).

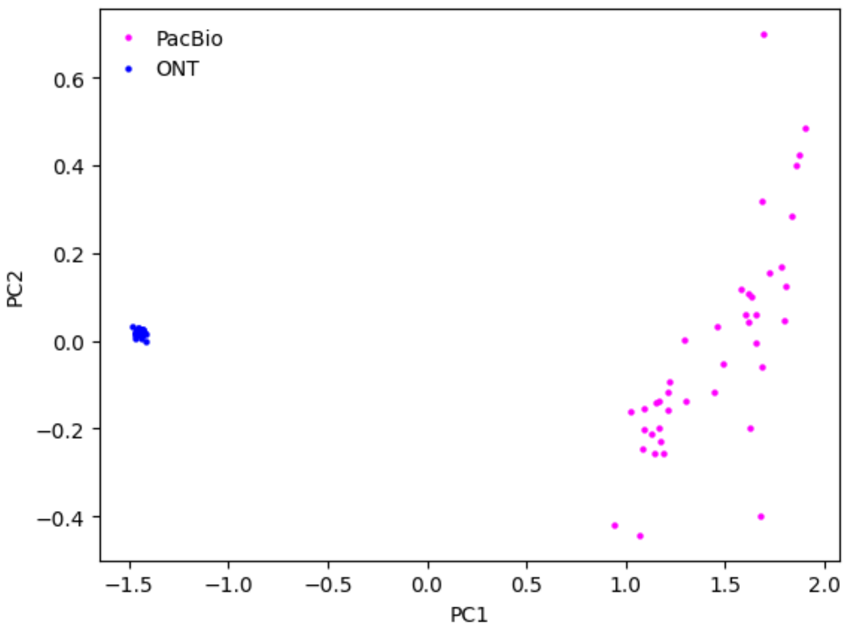


Supplementary Figure 8, PCA plot for CpG modification rate in 41 paired PacBio and ONT samples in AoU v8 cohort.

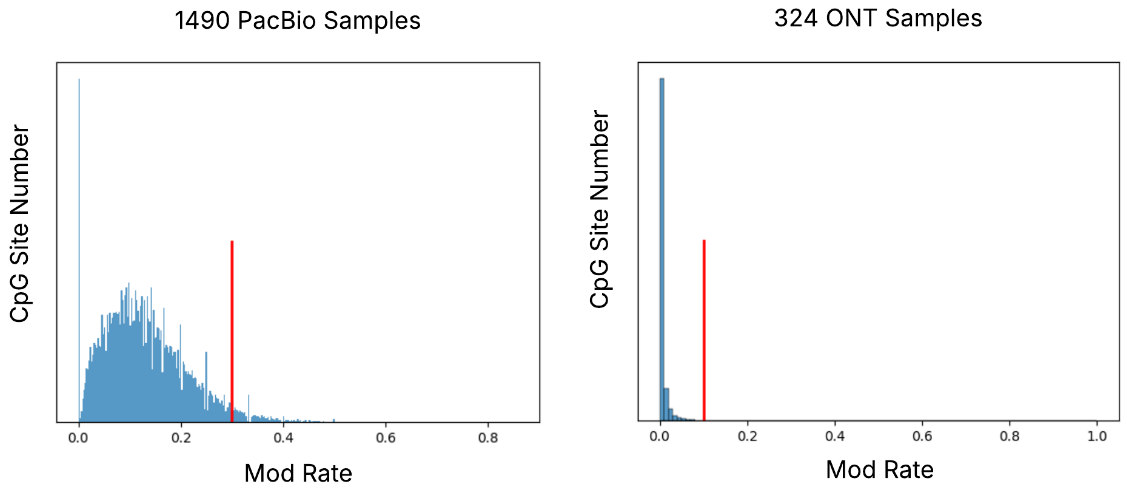

Supplementary Figure 9, The modification rate distribution.

Methylated count normalized by coverage of the CpG sites across all the 1490 PacBio long-read samples and 324 ONT samples. The red lines denote the threshold determining the 5mCpG is hyper-methylated.

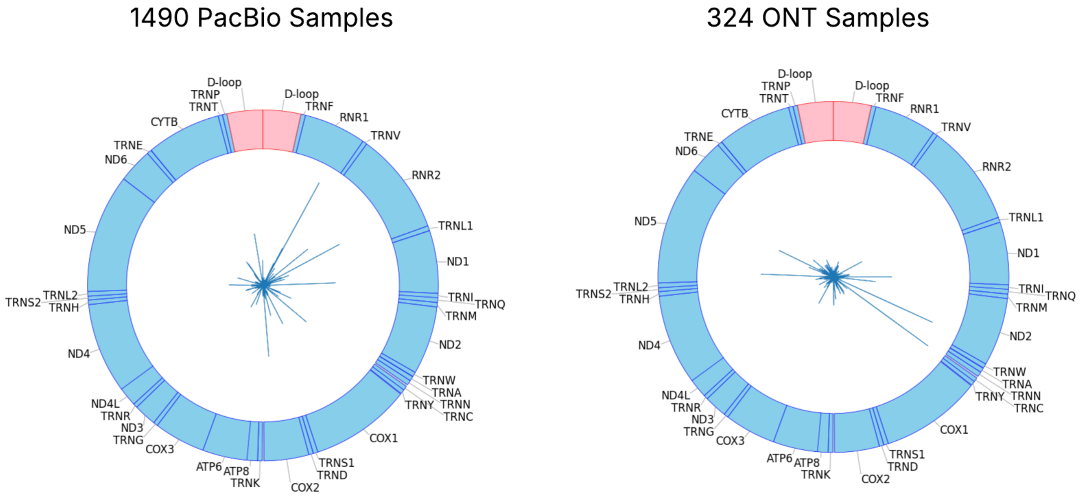


Supplementary Figure 10, Recurrent hypermethylated CpG sites across PacBio (modification rate threshold = 0.3) and ONT samples (modification rate threshold = 0.1).

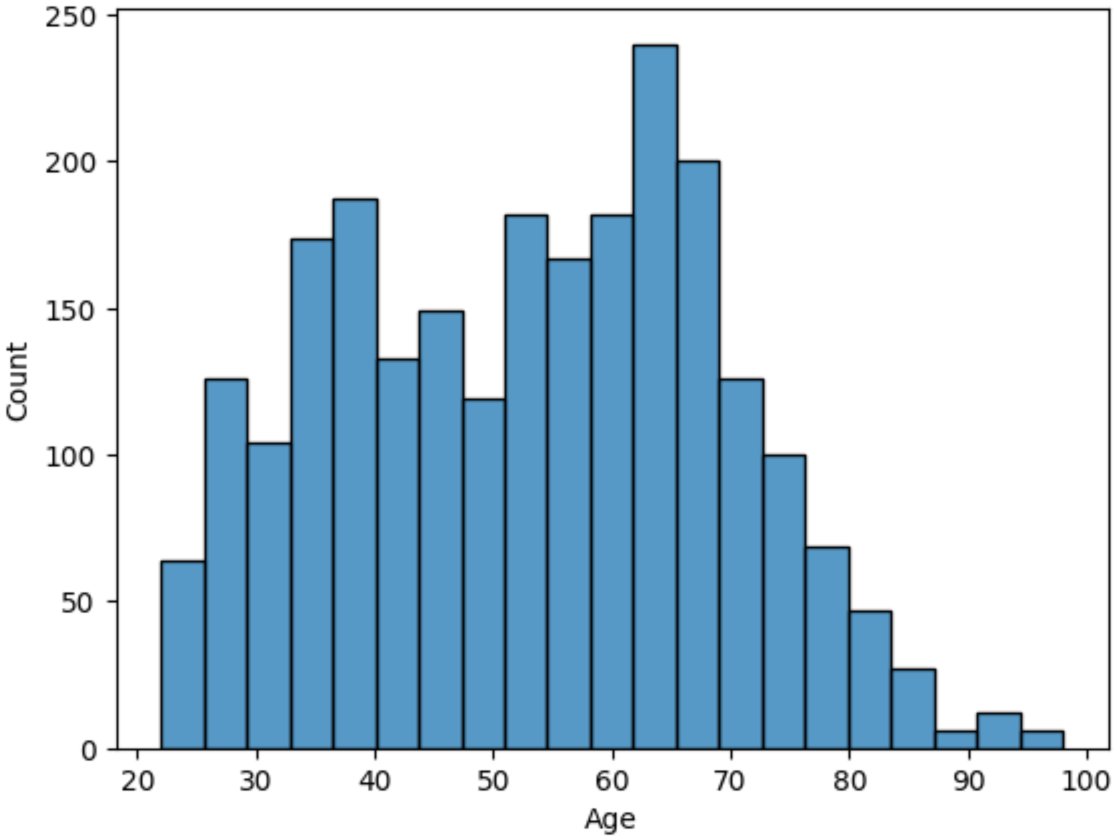


Supplementary Figure 11 Age distribution in AoU v8 release.

1014

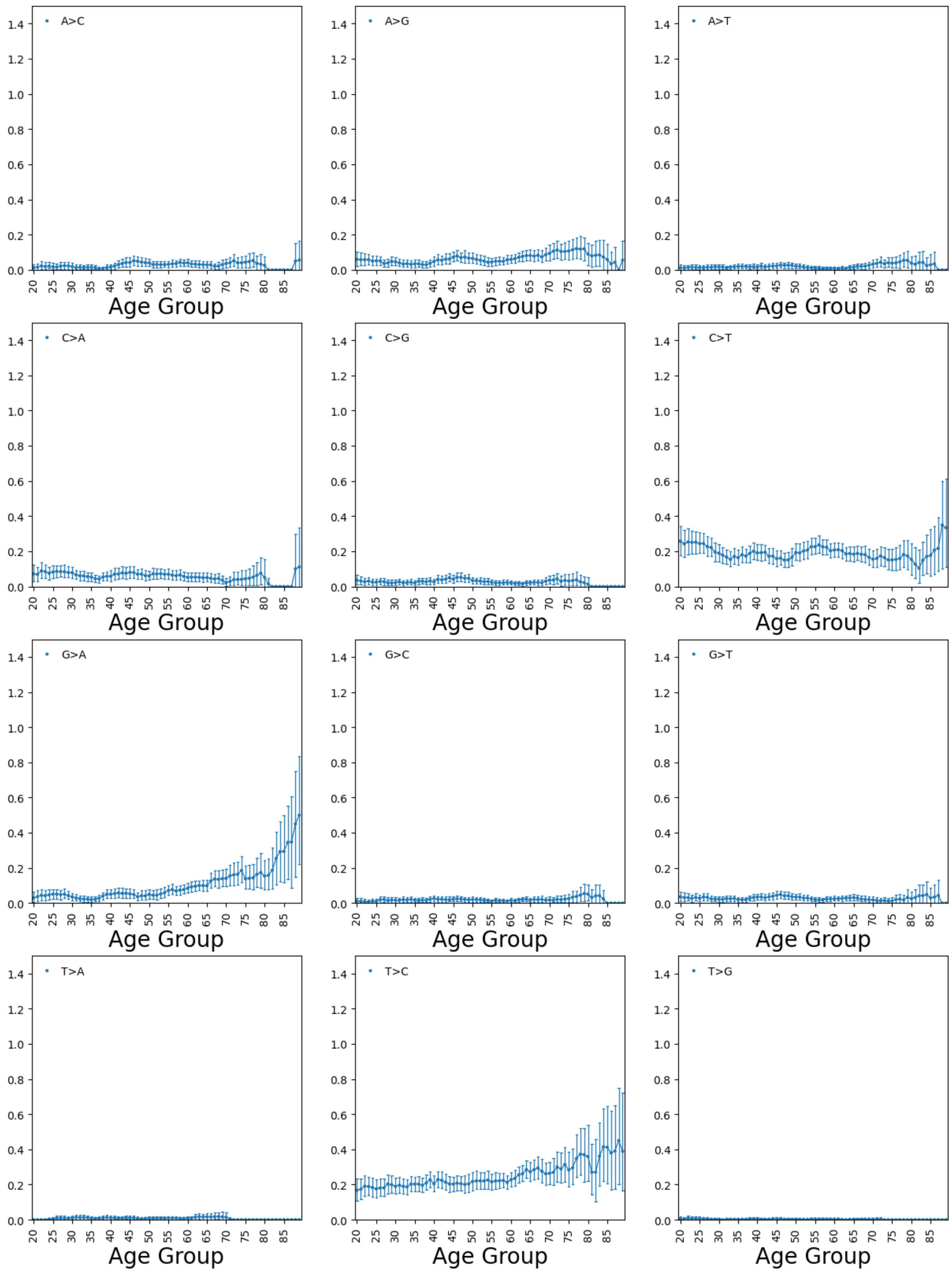

Supplementary Figure 12. The number of different variant types in Age groups (sliding window of 10).

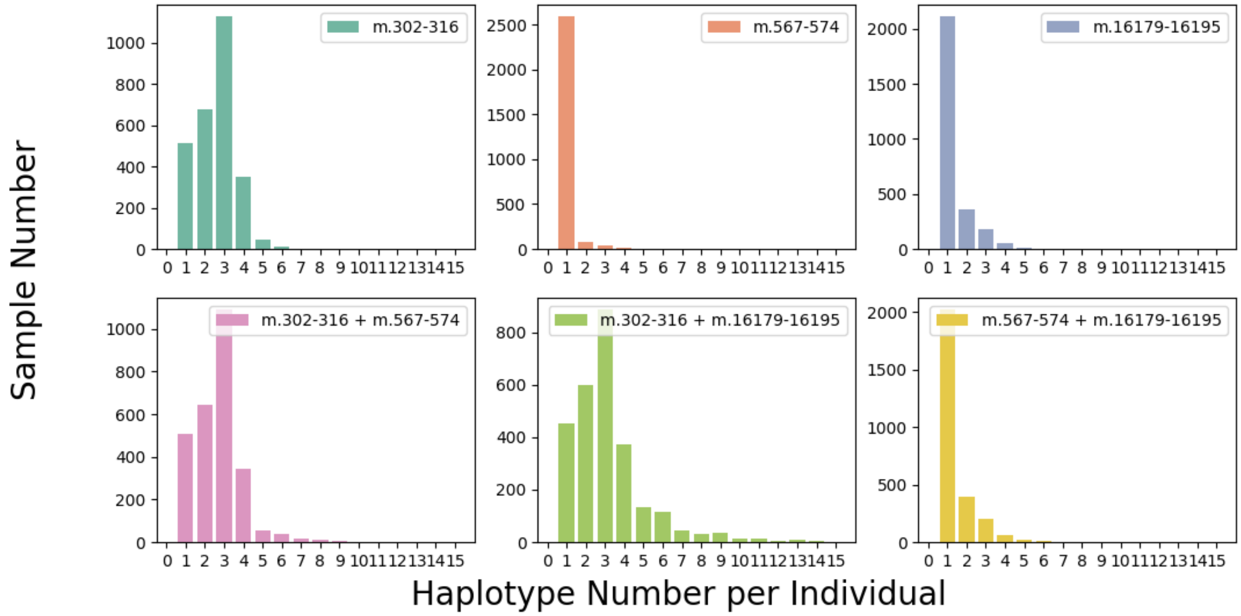


Supplementary Figure 13. Number of length heteroplasmic variants per individual and the joint alleles distribution.

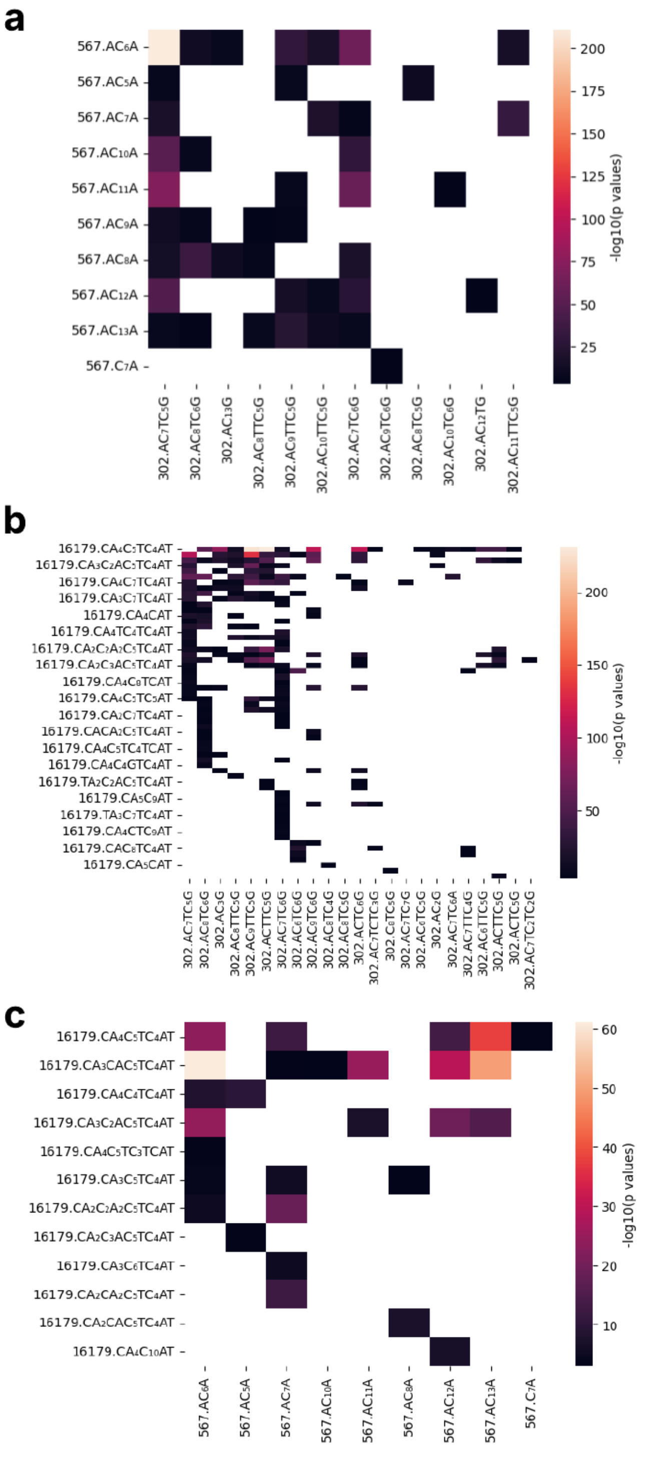
a
567.AC6A
567.AC5A
567.AC7A
567.AC10A
567.AC11A
567.AC9A
567.AC8A
567.AC12A
567.AC13A
567.C7A
302.AC7TC5G
302.AC8TC6G
302.AC13G
302.AC8TTC5G
302.AC9TTC5G
302.AC10TTC5G
302.AC7TC6G
302.AC9TC6G
302.AC8TC5G
302.AC10TC6G
302.AC12TG
302.AC11TTC5G
-log10(p values)
b
16179.CA4C5TC4AT
16179.CA3C2AC5TC4AT
16179.CA4C7TC4AT
16179.CA3C7TC4AT
16179.CA4CAT
16179.CA4TC4TC4AT
16179.CA2C2A2C5TC4AT
16179.CA2C3AC5TC4AT
16179.CA4C8TCAT
16179.CA4C5TC5AT
16179.CA2C7TC4AT
16179.CACA2C5TC4AT
16179.CA4C5TC4TCAT
16179.CA4C4GTC4AT
16179.TA2C2AC5TC4AT
16179.CA5C9AT
16179.TA3C7TC4AT
16179.CA4CTC9AT
16179.CAC8TC4AT
16179.CA5CAT
-log10(p values)
c
16179.CA4C5TC4AT
16179.CA3CAC5TC4AT
16179.CA4C4TC4AT
16179.CA3C2AC5TC4AT
16179.CA4C5TC3TCAT
16179.CA3C5TC4AT
16179.CA2C2A2C5TC4AT
16179.CA2C3AC5TC4AT
16179.CA3C6TC4AT
16179.CA2CA2C5TC4AT
16179.CA2CAC5TC4AT
16179.CA4C10AT
567.AC6A
567.AC5A
567.AC7A
567.AC10A
567.AC11A
567.AC8A
567.AC12A
567.AC13A
567.C7A
-log10(p values)

Supplementary Figure 14. Fisher’s Exact test for the association for length heteroplasmic alleles across 229985 reads.

Allele pairs with p-value < 0.001 are included in these plots.

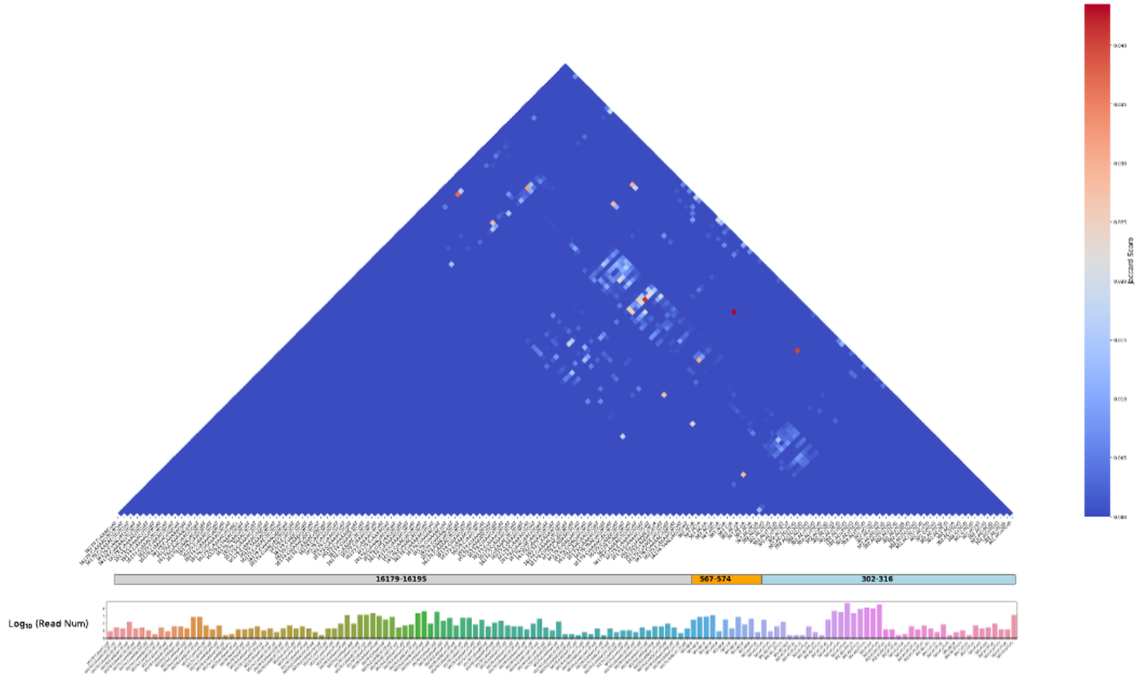


Supplementary Figure 15 Co-occurrence of length heteroplasmies.

We aggregated the binary matrix of variants versus reads in all the AoU v8 PacBio samples and test the co-occurrence of 3 length heteroplasmies in chrM:302-301, chrM:567-574, chrM:16179-16195. The Jaccard score is calculated by the number of reads supporting the co-occurrence of two length heteroplasmies, divided by the total number of reads supporting each length heteroplasmy. The bar plot shows the number of reads supporting each allele present in the heatmap.

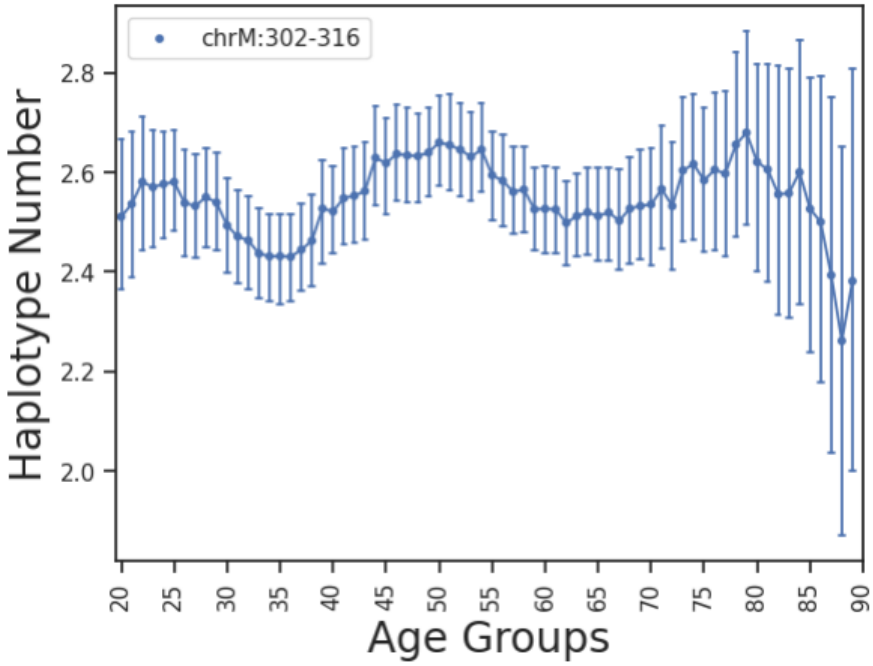


Supplementary Figure 16 Haplotype Number of chrM:302-316 in Age groups (sliding window of 10).

The haplotype is reconstructed by stitching the alternative alleles to the reference sequences according to the variant-read binary occurrence matrix of each sample.

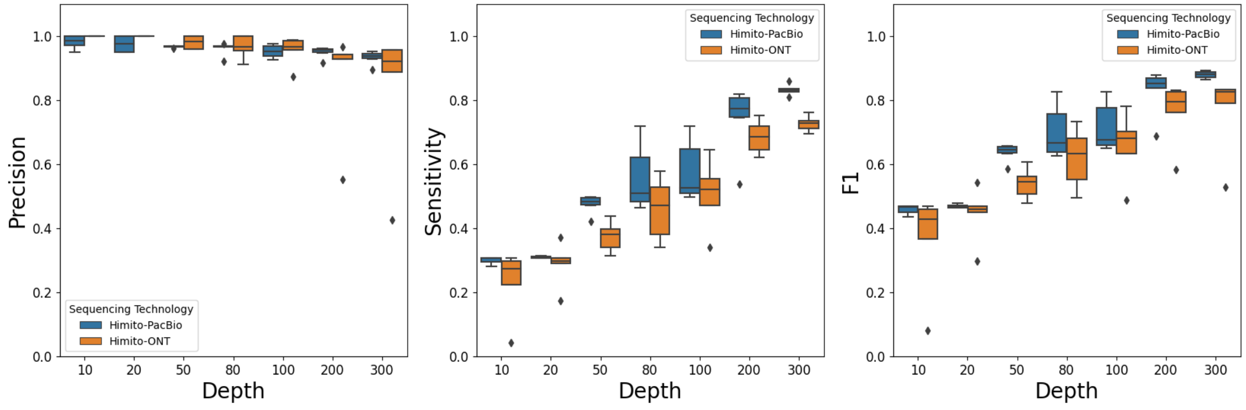


Supplementary Figure 17 HapMap mixture benchmarking using different long read sequencing technology (PacBio, ONT).
